# In vivo imaging in transgenic songbirds reveals superdiffusive neuron migration in the adult brain

**DOI:** 10.1101/2023.07.14.548876

**Authors:** Naomi R. Shvedov, Sina Analoui, Theresia Dafalias, Brooke L. Bedell, Timothy J. Gardner, Benjamin B. Scott

## Abstract

Neuron migration is a key phase of neurogenesis, critical for the assembly and function of neuronal circuits. In songbirds, this process continues throughout life, but how these newborn neurons disperse through the adult brain is unclear. We addressed this question using *in vivo* two-photon imaging in transgenic songbirds that express GFP in young neurons. In juvenile and adult birds, migratory cells were present at a high density, traveled in all directions, and made frequent course changes. Notably, these dynamic migration patterns were well fit by a superdiffusive model. Simulations revealed that these diffusion-like dynamics were sufficient to disperse new neurons throughout the song nucleus HVC. These results suggest that diffusion-like migration may underlie the formation and maintenance of nuclear brain structures in the postnatal brain and indicate that transgenic songbirds are a useful resource for future studies into the mechanisms of adult neurogenesis.

**Highlights:**

- Transgenic songbirds express GFP in a neurogenic lineage
- GFP expression is strong and sparse enough to track single cells *in vivo*
- Adult neuron migration is well fit by a superdiffusive model
- Superdiffusive migration is sufficient to populate HVC in simulation

## Introduction

In the songbird, new neurons are continually added to forebrain circuits throughout life. These new neurons are added to a variety of regions, including HVC, a nucleus in the song circuit, which is involved in learned vocal communication^1^. In HVC, the addition of new projection neurons and interneurons are thought to play an important role in behavioral plasticity and tissue resilience^2–6^. These new neurons are born from neural stem cells in the ventricular zone (VZ) and migrate hundreds of micrometers to reach their integration targets in HVC and beyond^4,7^. How these new neurons migrate through the adult brain is unknown. Although neuron migration has been well studied in the vertebrate forebrain during development, the adult brain presents a different set of challenges to migration and integration, and may involve new mechanisms^8^.

Previous histological and *in vitro* experiments in adult songbirds described the close association of some young neurons with “radial glia” guide cells, similar to migration in the developing mammalian cortex^1,7,9^. However, many areas of the songbird brain that receive new neurons, including the song nucleus HVC, contain relatively few radial fibers suggesting the possibility of other migratory mechanisms^10^. Retroviral labeling studies have revealed that migratory neurons in juvenile zebra finches follow tortuous paths that resemble a random walk, and do not appear to exclusively follow radial fibers or blood vessels^11^. Together, these observations suggest the existence of a non-radial form of migration in the postnatal songbird brain. However, a detailed description of this process and whether it continues into adulthood remains unknown.

To address this question, we sought to quantitatively characterize the cellular dynamics of newborn neurons as they migrated through the brains of juvenile and adult zebra finches. We took advantage of a line of transgenic zebra finches that express green fluorescent protein (GFP) under the ubiquitin C (UBC) promoter^12^. The UBC-GFP transgenic strain was produced to demonstrate the efficiency of lentiviral based transgenesis in songbirds, but GFP expression had not been characterized in detail. Here we show that GFP is strongly expressed in a neurogenic lineage that includes both young and mature neurons and that expression is sparse enough and bright enough to enable single cell identification and tracking *in vivo*. We used timelapse two photon microscopy (2PM) to follow populations of GFP-positive (GFP+) cells for up to 12 hours as they migrated through the brain and applied a three-dimensional (3D) manual tracking pipeline to quantify the migratory dynamics of over 800 migratory neuroblasts.

We found that migratory cells were present in high density within the juvenile and adult songbird brain. Migratory neurons were not confined to a particular region or pathway, but were found throughout the hyperpallium above HVC, and the nidopallium in and around HVC, including in the brains of female birds. New neurons moved in different directions and made frequent turns, and their trajectories were well fit by a superdiffusive model. Finally, using statistics measured from *in vivo* data, we developed a quantitative simulation of the migration. This simulation revealed that this diffusive process was sufficient to populate HVC within 3 weeks, the biologically relevant timescale scale for migration in songbirds^7^. Together these results demonstrate a dynamic form of neuron migration that may allow for flexible and efficient dispersal through the postnatal brain.

## Results

### UBC-GFP transgenic songbirds express GFP in a neurogenic lineage

We first characterized GFP expression in the brains of UBC-GFP zebra finches in histological sections. We found that GFP was expressed throughout the forebrain, including the song nuclei, but varied across regions and cell types (**Figures 1**, **S1**). Most GFP+ cells (90.8%, 188/207 cells, n = 1 bird) expressed Hu, a neuronal lineage marker (**Figure 1D**), and the majority (64.2%, 258/402 cells, n = 4 birds) were positive for NeuN, a marker for mature, post-migratory neurons (**Figure 1B**).

**Figure 1.**
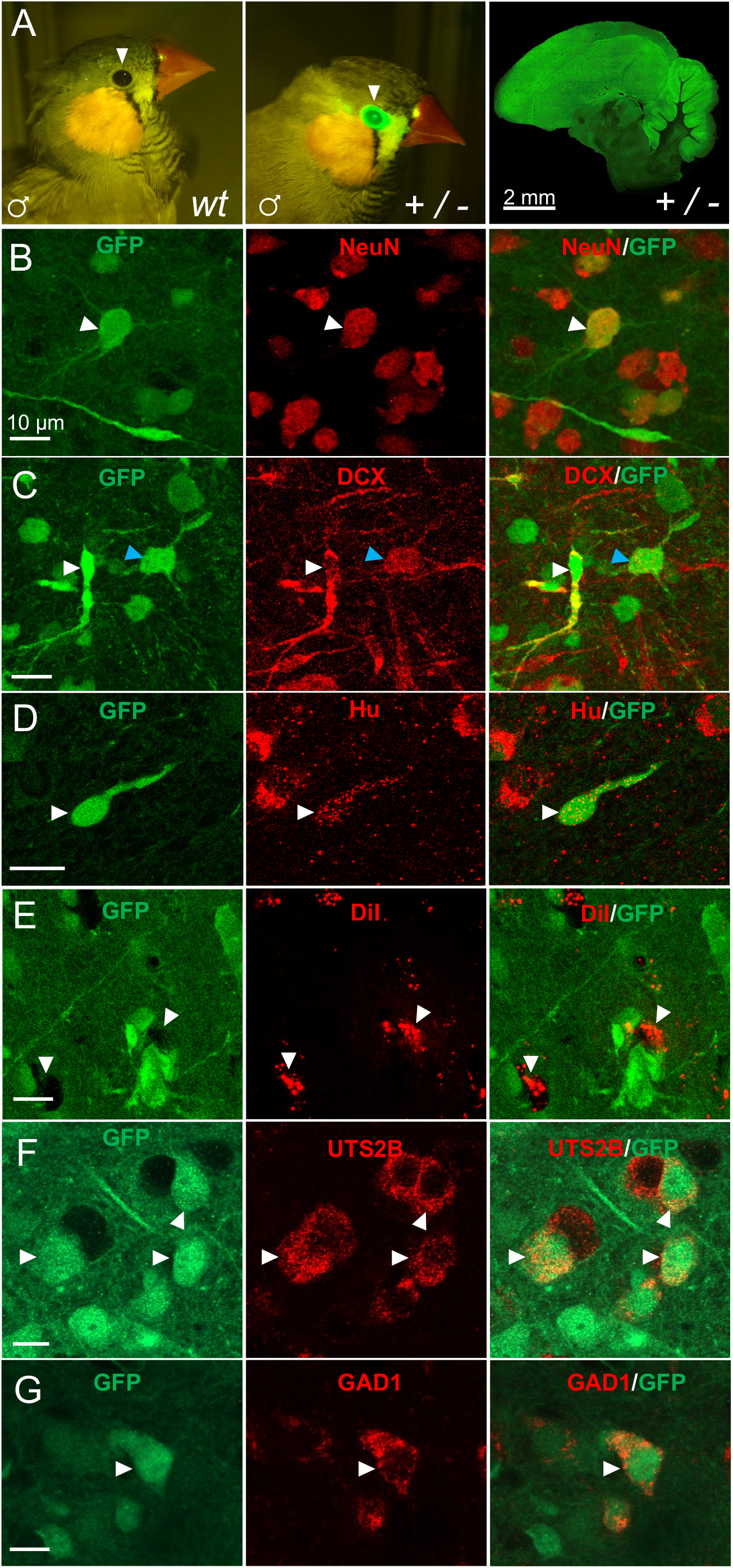
UBC-GFP transgenic songbirds express GFP in a neurogenic lineage. A. *Left:* Adult male wildtype finch under blue light excitation. No GFP fluorescence is observed (white arrow); *Middle:* Adult male transgenic finch under blue light excitation. Bright GFP fluorescence is observed around the eye (white arrow); *Right:* GFP expression found throughout transgenic male adult brain. B. Mature neuron co-labeled with GFP and NeuN (white arrow). C. GFP+ cell with bipolar neuron morphology labeled with DCX (white arrow). Cell with multipolar morphology labeled with GFP and DCX (blue arrow). D. GFP+ bipolar cell co-labeled with Hu (white arrow). E. GFP-negative, DiI-labeled HVC_X_ projectors (white arrows). F. HCR-FISH for UTS2B (red) and IHC staining for GFP (green). GFP+/UTS2B+ (HVC_RA_ neurons) shown with white arrows. G. HCR-FISH for GAD1 (red) and IHC staining for GFP (green). GFP+/GAD1+ (mature interneurons) shown with white arrow.

In song nucleus HVC, we evaluated GFP expression in three well-studied neuronal subtypes: HVC to RA glutamatergic projection neurons (HVC_RA_), HVC to Area X glutamatergic projection neurons (HVC_X_), and GABAergic interneurons (HVC_INT_). In HVC_X_ projection neurons that were retrogradely labeled from Area X with DiI, almost no GFP+ cells were found (n = 2/141 cells) (**Figure 1E**). In HVC_RA_ projection neurons, identified using in-situ hybridization (ISH) for Urotensin 2B (UTS2B), more than half of the cells (60.8%, n = 192/316 cells) were GFP+ (**Figure 1F**). In HVC_INT_, identified using ISH for glutamate decarboxylase (GAD1), half were GFP+ (49%, n = 98/200 cells) (**Figure 1G**).

In addition to mature neuron types, GFP was also expressed in doublecortin (DCX) positive putative migratory neurons (45.7% of DCX+ cells were GFP+; n = 288/630, n = 4 birds). Although the density of labeling made complete morphological characterization difficult to achieve for all cells, many GFP+/DCX+ cells exhibited a bipolar morphology with a small, elongated soma, or were multipolar with a larger soma (**Figure 1C)**, indicative of migratory neurons at different stages of maturity^13^. As a proportion of GFP+ cells, GFP+/DCX+ were relatively abundant; 41% of GFP+ cells expressed DCX (174/424 cells, n = 4 birds). Together this data indicates that in the HVC and surrounding tissue of UBC-GFP transgenics, GFP expression appears to be primarily expressed in cells of a neurogenic lineage, including a high density of putative migratory neurons.

### Transplanted GFP+ cells migrate and become neurons in host brains

To further study the identity of GFP+ cells in UBC-GFP birds, we transplanted a small piece of living brain tissue (5-10 mg) from transgenics into a wildtype hosts and then evaluated GFP expression in the host brain at different times post-surgery (n = 6 birds; **Figure 2A**, **Table S1**). Three weeks after transplantation, GFP+ cells were found in the host brain up to 350 μm from the edge of the graft. These cells exhibited a bipolar morphology, characteristic of migratory neurons (**Figure 2B**, n = 21 cells, 1 bird). Five weeks after transplantation, GFP+ cells were observed up to 2 mm from the graft and were commonly multipolar (average distance from the graft: 1042 μm +/- 626.23 μm; n = 26 cells from 2 birds; **Figure 2C**). Seven to nine weeks post transplantation, GFP+ cells which had dispersed into the host tissue now exhibited mature neuronal morphology, had spiny dendrites, and expressed NeuN (n = 9/9 cells, 3 birds; **Figure 2D-G**). These experiments indicate that the brains of UBC-GFP zebra finches contain GFP+ cells that are capable of migrating into the brain of another animal and differentiating into mature neurons.

**Figure 2.**
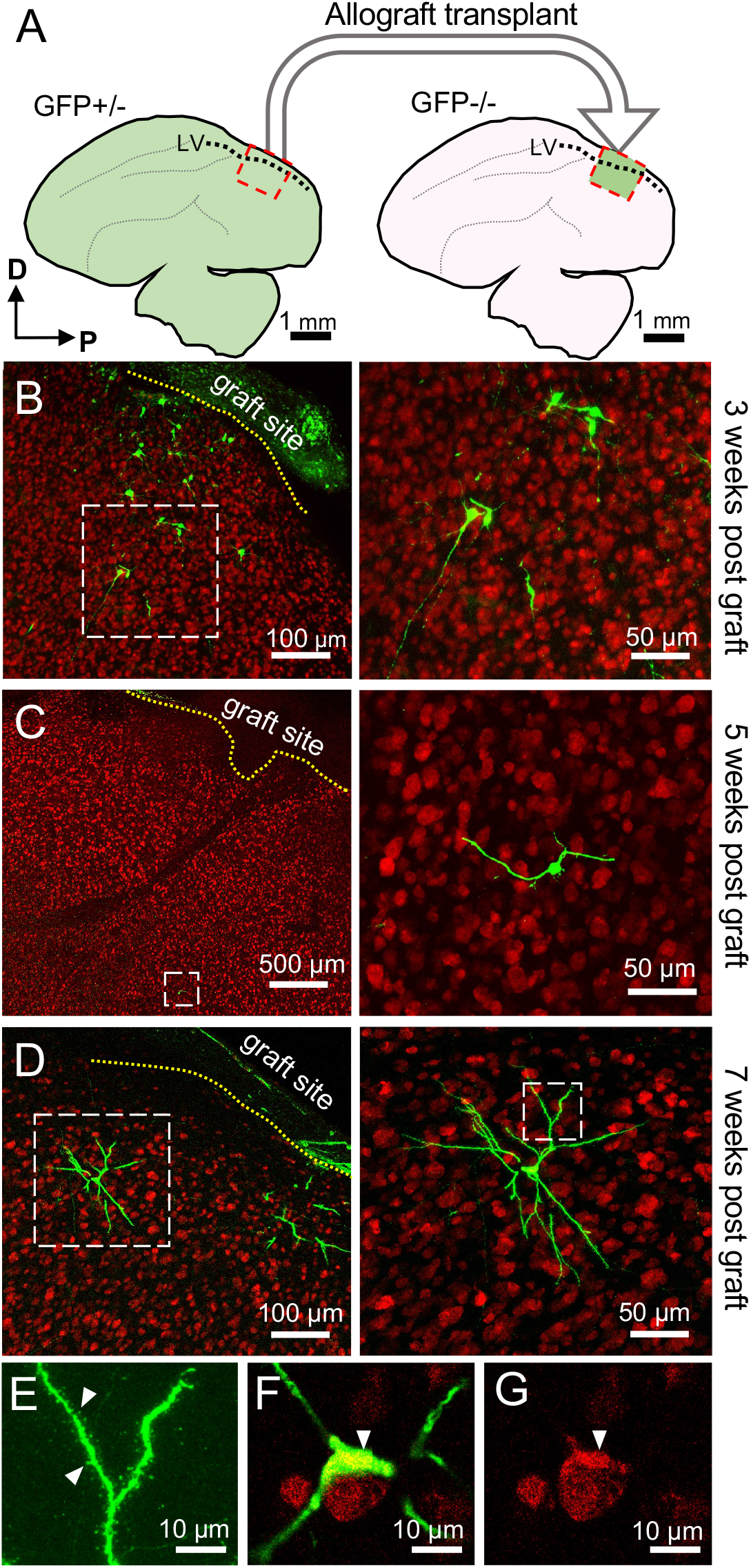
GFP+ cells from donor transplants migrate into host brains and become neurons. A. Schematic of graft procedure. B. Histological section from the host brain three weeks after transplantation showing GFP+ cells (green) and NeuN+ neurons (red); *Left:* Yellow dotted line indicates the graft-host interface; *Right:* Zoom in on the white dashed region shown on right showing GFP+ migrating cells. C. *Left:* histological section showing GFP+ cell located approximately 1.5 millimeters from the graft boundary (dotted yellow line); *Right:* Zoom in of white dashed region shown on left. D. *Left:* histological section from the host brain six weeks after transplantation. GFP+ mature neurons (green) and mature NeuN+ neurons (red). Yellow dotted line is the graft-host interface; *Right*: Zoom in on the white dashed region shown on left showing a GFP+ neuron. E. Zoom in of dashed region in right panel of D. White arrows point to dendritic spines. F. Zoom in of GFP+ cell shown in D and E. White arrow points to soma. G. White arrowhead points to NeuN+ soma of cell shown in F.

### *in vivo* imaging in UBC-GFP songbirds reveals dynamic cell migration

We next sought to determine if GFP expression was bright and sparse enough to allow for imaging of migrating cells *in vivo*. We implanted UBC-GFP birds with a 3 mm cranial window and kinematic headplate^14^ and performed 2P time lapse volumetric microscopy (**Figure 3A-C**). Using this approach, we could resolve large numbers of individual cells down to 350 μm below the brain’s surface. Many cells exhibiting the bipolar morphology characteristic of migratory neurons were found both above the VZ in the hyperpallium and below the VZ in regions of the nidopallium, including HVC (**Figure 3B**, **Video S1**). Timelapse volumetric imaging through implanted cranial windows revealed a high density of GFP+ migratory cells throughout these regions that were moving in different directions and making frequent turns (**Video S2**). To enable imaging of cells in deeper areas, we used a microprism approach in which a 1.5 mm microprism was bonded to a cranial window with optical adhesive and surgically implanted^15–17^ (**Figure 3B**, see **Methods**). This microprism approach allowed imaging of cells up to 800 μm below the brain surface (**Figure S3A,B**). Using a combination of both cranial window and microprism approaches, we imaged migration in 8 birds (5 males, 3 females) for between 3 - 12 hours and manually tracked 876 trajectories across all volumes (109.5 +/- 44.6 trajectories per animal; **Figure 3D-F, Table S2**).

**Figure 3.**
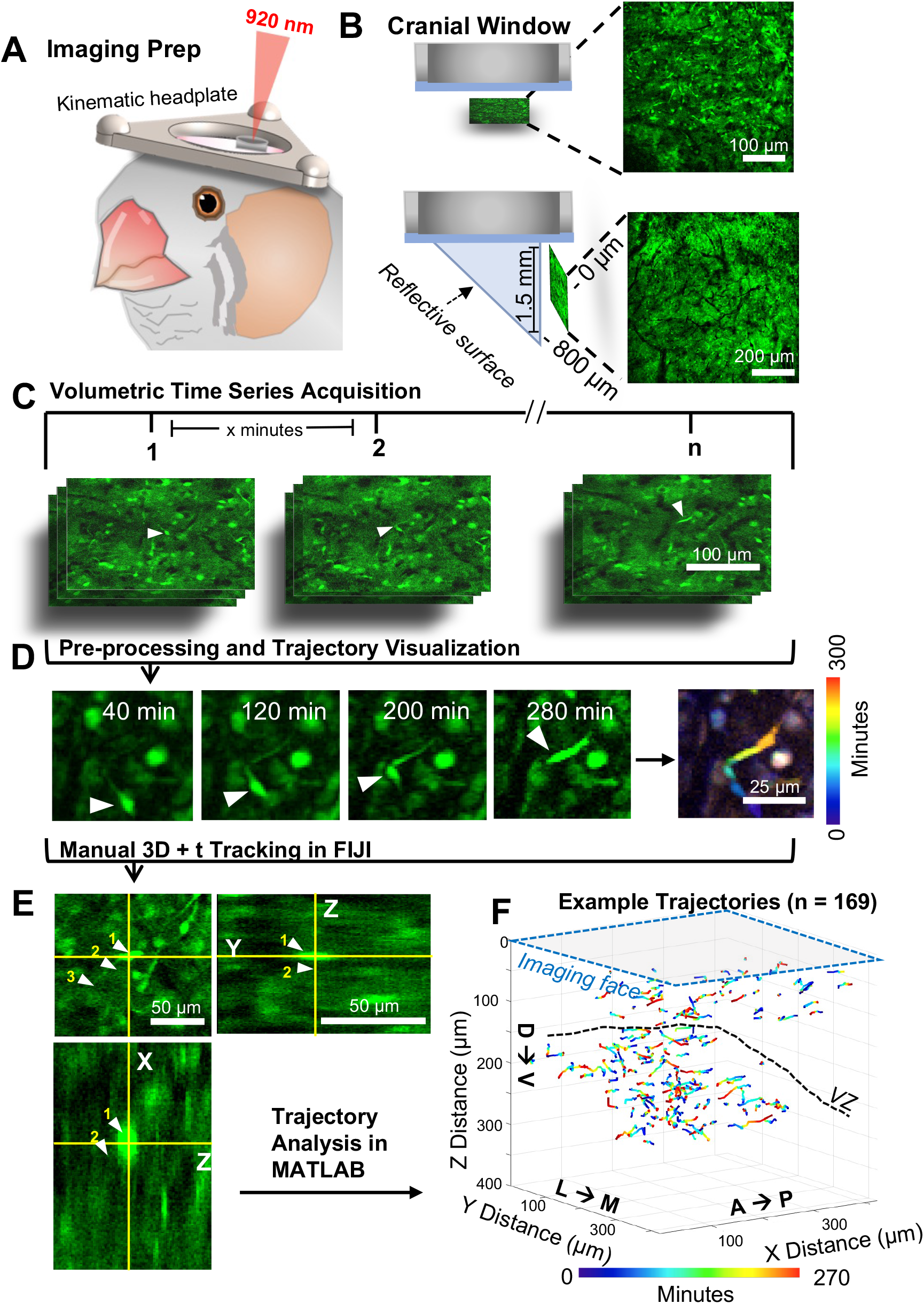
Migratory GFP+ cells can be imaged in UBC-GFP birds using two-photon microscopy. A. Imaging preparation. Kinematic headplate and optical implant shown. 920 nm light beam shown entering optical implant. B. *Top:* Cranial window implant schematic and example FOV; *Bottom:* microprism implant schematic and example FOV. C. Design for 2-photon volumetric time-lapse recordings. D. Visualization of a single migrating cell (white arrow) over time. E. Schematic of manual tracking in 3D over time, performed in FIJI using a custom plug-in, for a single cell. White arrows and yellow numbers indicate positions of the cell’s soma at future timepoints. F. Example tracked trajectories, temporally color-coded from a single juvenile animal both above and below the VZ.

We first evaluated dynamics of migratory cells in juvenile HVC (9 - 13 min per volume, 425 µm x 425 µm x 310 µm, n = 143 cells, 2 birds; **Figure 4**). Migrating cells in HVC were dispersed throughout the imaging field of view (FOV) and moved in all directions (**Figure 4C-E**). We did not detect a bias for a particular direction in the horizontal plane (Rayleigh’s test, p = 0.83, n = 143 cells, 2 birds; **Figure 4F**). These cells also made frequent turns (mean tortuosity = 1.83 +/- 0.69; **Figure 4H**) and moved along their path at speeds ranging from 7.0 to 45 μm/h (mean 14.65 +/- 5.29 μm/h; **Figure 4G**). These dynamics were similar to those of retrovirally labeled migratory neurons in the juvenile songbird HVC^11^.

**Figure 4.**
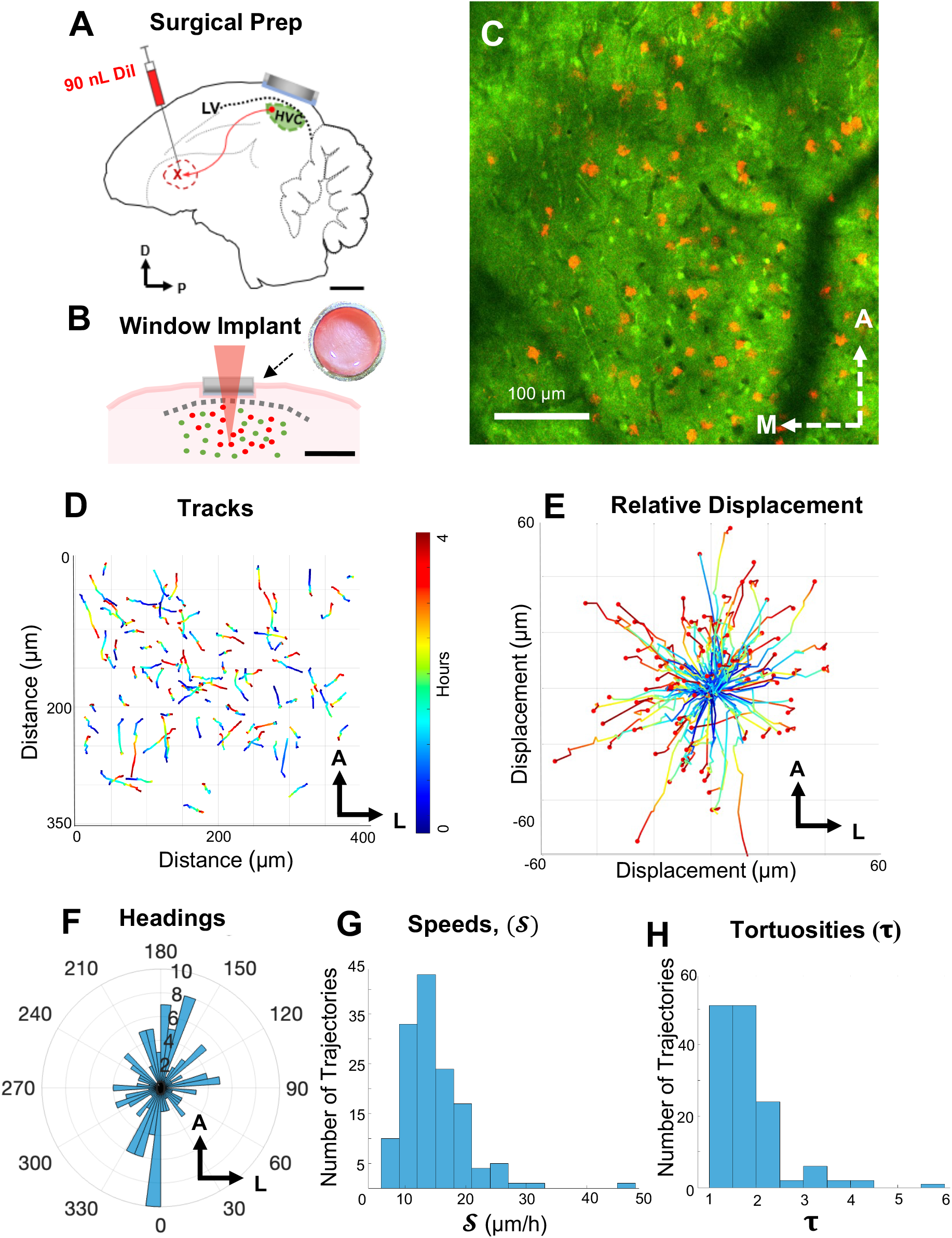
Dynamics of neuroblast migration in male zebra finch HVC. A. Surgical preparation of zebra finch for HVC imaging, retrograde injection of DiI into Area X and cranial window placement over HVC coordinates. B. Schematic of cranial window implant with 2P stimulation (920 nm) of DiI+ cells shown in red. C. 2P microscopy image of a single plane in HVC showing GFP expression (green) and DiI+ HVC_X_ projecting cells (red). D. Tracked trajectories of 96 migrating cells. Color indicates the time within the 4.5 hour duration imaging session. E. Relative displacement of cells shown in D with normalized start positions. F. Histogram of cell heading angles in the horizontal plane (Rayleigh’s test, p = 0.83; n = 143 cells, n = 2 birds). G. Distribution of average speeds of migrating cells (n = 143 trajectories, n = 2 birds). Mean speed = 14.65 +/- 5.29 µm/h. H. Distribution of tortuosities of migrating cells (n = 143 trajectories, n = 2 birds). Mean tortuosity = 1.83 +/- 0.69.

Similar dynamics were observed in the brains of adult birds (86-800+ dph, 3 males, 3 females, n = 594 cells). Both within HVC and in other regions of the nidopallium, we observed a large number of migrating cells that were distributed throughout the tissue and dispersing in all directions while making turns (**Video S2**). Interestingly, cells in superficial regions (within 400 microns of the surface) exhibited a slight but statistically significant ventral bias (1.22 μm/h ventrally; t-test, p < 0.005; n = 115 cells, 3 birds; **Figure S3C-E**) while deeper cells (over 400 microns from the surface) exhibited no ventral bias (t-test, p = 0.134; n = 133 cells, 3 birds). In addition, deeper cells had a significantly higher tortuosity than superficial cells (two-sample t-test, p < 0.0005; **Figure S3G**).

Together, these results suggest the presence of a high density of migratory neurons in the brains of juvenile, adult, male and female zebra finches.

### Population analysis reveals an autonomous, disordered migration process

We next wondered whether the migration dynamics of nearby cells were synchronized or whether cells moved in an independent fashion. We evaluated the correlation in heading direction and speed between 26,642 pairs of cells across 4 birds (105 +/- 44 simultaneously recorded cells per bird). We did not observe a significant correlation in heading directions between pairs of cells (**Figure 5A-B**) (2-sample Kolmogorov-Smirnov test, p = 0.12). However, we did observe a small but significant correlation in the speed of cell migration between pairs of cells (p > 0.001) (**Figure 5C**). This correlation in speed was slightly inversely correlated with the distance between the pairs of cells (**Figure 5D**).

**Figure 5:**
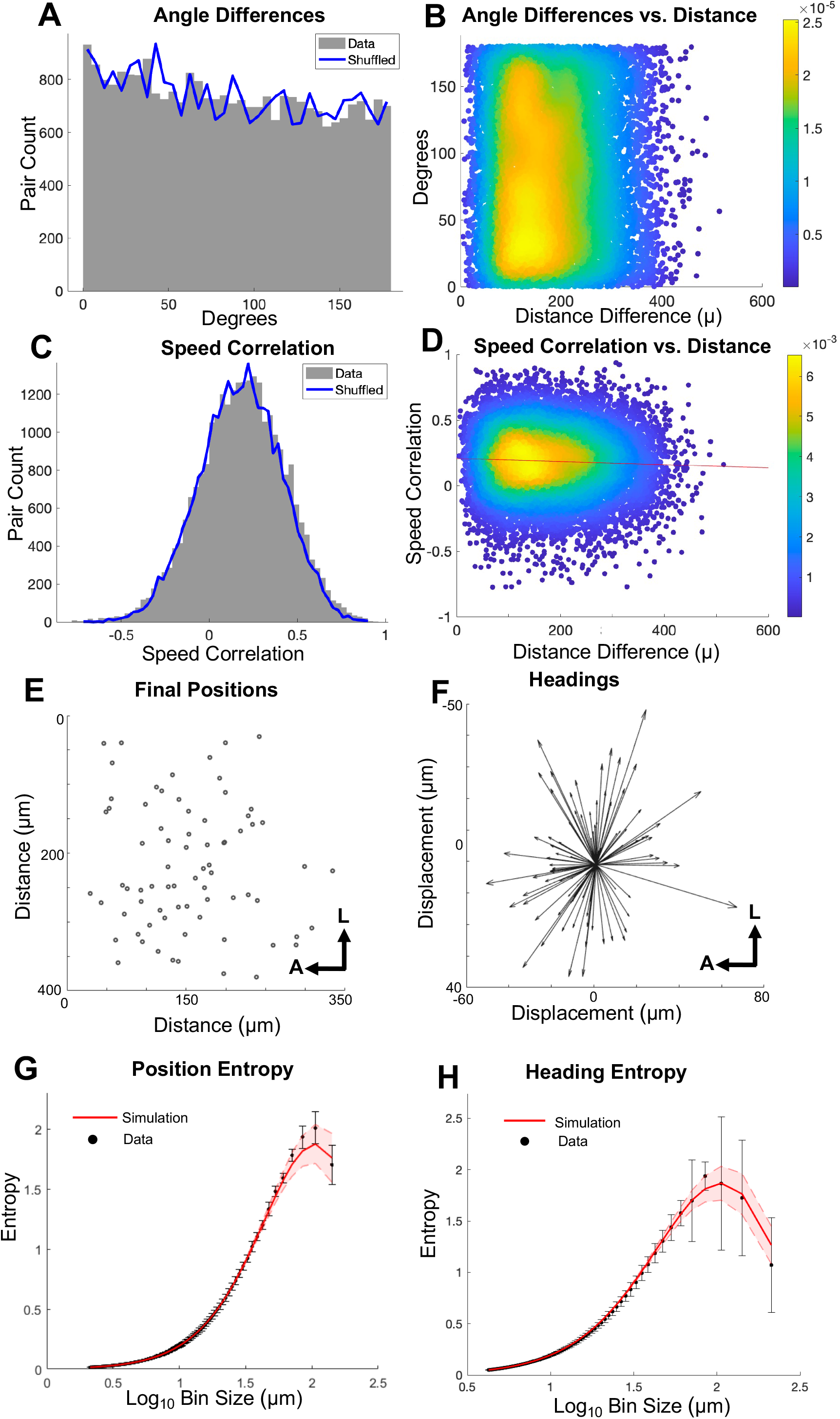
Population analysis reveals an autonomous, disordered migration process. A. Distribution of angle differences between the trajectories of 26,642 pairs of cells (gray) from 4 birds (3 males, 1 female) compared to shuffled data (blue line) (2-sample Kolmogorov-Smirnov test, p = 0.12). B. Scatterplot of pairwise angle differences and distance difference between the cell pair. Each point is a cell pair, color indicates kernel density estimate. No significant relationship between angle difference and distance between pairs (linear regression, p = 0.9). C. Distribution of speed correlations (Spearman’s) between the trajectories of 26,642 pairs from 4 birds (3 males, 1 female) compared to shuffled data pairwise speed correlations (2-sample Kolmogorov-Smirnov test, p < 0.0005). D. A small but significant relationship found between speed correlation and distance between pairs (linear regression, slope = -0.000118 change in correlation value per micron, p < 0.0005, R^2^ = 0.0018). This effect was slightly greater than was expected by shuffled data (linear regression, slope = -8.0685e-05 change in correlation value per micron, p < 0.0005, R^2^ = 0.000935). E. Positions of 96 simultaneously recorded migratory cells in the imaging volume, projected into a single plane. F. Headings of 96 simultaneously recorded migratory cells in the imaging volume. G. Entropy average of observed cell positions across time (black dots) plotted with mean of 1000 simulations of maximum entropy model run with mean number of position observations across time (red line). Black error bars are the standard deviations of entropies of positions across 18 timepoints within 4.5 hours. Shaded red region is the standard deviation of entropy across 1000 simulations. H. Entropy of trajectory directions across a 4.5 hour imaging session (black dots) plotted with mean of 1000 simulations of maximum entropy model (red line).

Next, to detect and evaluate the presence of ordered patterns in the movements of cells, we estimated the Shannon entropy of migration dynamics (**Figure 5E,F**, see **Supplemental Methods**). We found that the entropy of the position and heading of cells were well-fit by a uniform random distribution maximum entropy model across a range of spatial scales (**Figure 5G,H**). Together these results suggest that migration in the postnatal zebra finch forebrain is consistent with a disordered process in which cells show little correlation in their positions, heading and speed.

### Migrating neurons may flexibly use vascular scaffolds

We investigated if the migrating neuroblasts were associated with radial glia or vasculature during their migration, as described in previous work^7,18–23^. To evaluate association with radial glia, we counted instances of putative migratory neurons being in contact with vimentin positive (Vim+) fibers in histological sections. Consistent with previous work, only a subset of cells with migratory morphology were associated with Vim+ fibers in HVC^11^ and surrounding nidopallium (4/22 cells, n = 1 bird). This suggests that migrating neurons may not be dependent on Vim+ fibers for migration.

To evaluate association with vasculature, we labeled the vasculature with SR-101 at the end of *in vivo* imaging sessions and compared the migratory trajectories with the pattern of blood vessels (n = 115 cells, n = 2 birds). Over a 3-hour imaging session, 21% of cells exhibited trajectories that closely aligned to the path of the vasculature (n = 17/81 cells, 1 bird; See **Methods**). Interestingly, we observed that most vasculature-associated cells (64.7%, n = 11/17 cells) did not follow the path of the vessel for the entirety of their trajectories. Instead, they appeared to hop on and off the vasculature for a proportion of their trajectory. These cells followed vasculature for an average of 35.17 +/- 21.02 μm out of their total 48.64 +/- 17.06 μm long trajectories (71.8% of total path traveled spent on vasculature, on average). Cells that followed vasculature at some point during their trajectory were significantly faster on average than cells that were not associated with vasculature at all (15.55 +/- 17.90 vs 10.81 +/- 13.36 μm/h) (two-sample t-test, p < 0.001).

Similar results were observed across a longer imaging session (11.43 hours, 34 cells, 1 bird). The paths of a small number of cells (23.5%, 8/34) aligned to the paths of the SR-101-labeled vasculature for some segment of their trajectory (n = 8/34 tracks). Nearly all of the migratory cells that followed along vasculature paths did so transiently within the imaging session (87.5%, 7/8 trajectories, **Figure S4**). These cells followed vasculature for an average of 55.66 +/- 44.93 μm out of their total 95.39 +/- 47.58 μm long trajectories (∼52.7% of total path). Cells that followed vasculature at some point during their trajectory were significantly faster on average than cells that were not associated with vasculature at all (mean instantaneous velocity: 8.97 +/- 8.53 μm/h vs 6.36 +/- 6.06 μm/h; two-sample t-test, p < 0.01). These results suggest that migrating neurons in the postnatal zebra finch brain may flexibly use vasculature for a portion of their migration path.

### Neuron migration is well described by a superdiffusive model

Our observations suggest that new neurons in the postnatal songbird brain disperse in various directions, make frequent turns, and are largely uncorrelated in their migration patterns. Therefore we wondered whether their migratory behavior could be described by a diffusive model, similar to those applied to other migratory cell types, such as bacteria, mammalian T-cells, and ‘super-spreading’ cancer cells^24–26^. Quantitative analysis of diffusion in a system can offer insight into the biological mechanisms governing the migration of cell populations^27,28^. To evaluate a diffusive model of migration we computed the mean squared displacement (MSD) of the soma centroid over time from tracked cells (**Figure 6**). MSD for a single trajectory was computed using the following formula:

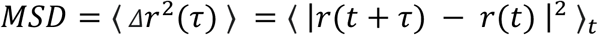

where *r*(*t*) is the soma position at time *t* and *r*(*t* + *τ*) was the new soma position at time *t* + *τ*^29^.

**Figure 6.**
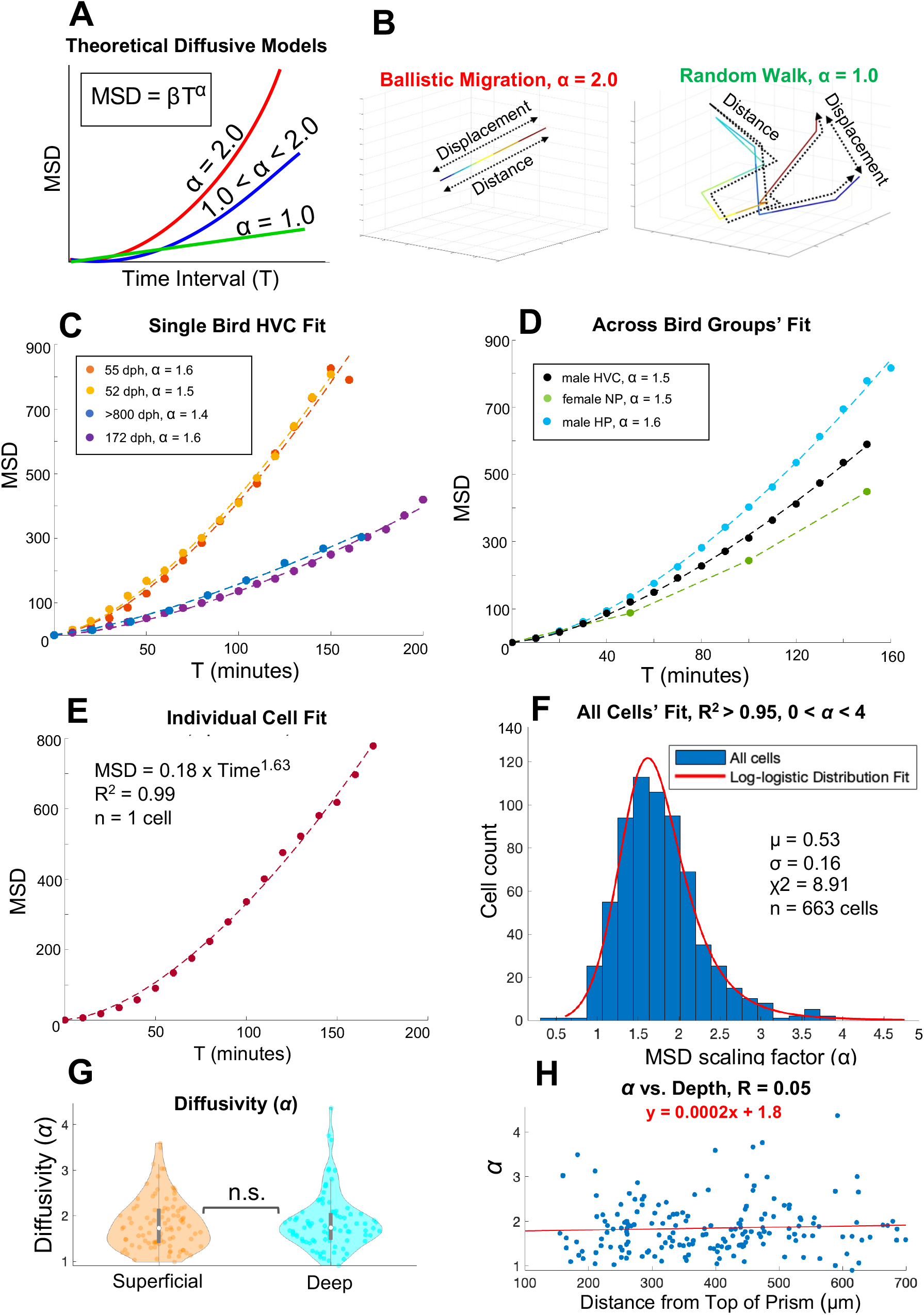
Migration dynamics are well-fit by a superdiffusive model. A. Schematic indicating MSD vs time (T) power law model and relevant *α* parameters under different diffusion types: ballistic (red), superdiffusive (blue), and random walk (green). B. *Left:* Schematic of ballistic migration; *Right:* schematic of random walk migration. C. Diffusion model fit for the average 2D MSD of 301 migratory cells from four male HVC’s (52 dph (n = 44), 55 dph (n = 95), 172 dph (n = 79), >800 dph (n = 83)). D. Model fits to the average 2D MSD of cells in different brain regions. Male HVC data includes 218 cells from 3 males, female nidopallium (NP) data includes 247 cells from 2 female birds, and male hyperpallium (HP) includes 143 cells from 2 males. E. Model fit to an example cells’ MSD. F. Histogram of the *α* parameter distribution across all cells (n = 663/940 cells with R^2^ > 0.95, *α* < 4). The data is well fit by a unimodal log-logistic distribution (red line). The average alpha is 1.7664 +/- 0.5004. G. Violin plots of MSD coefficient, *α,* from superficial (n = 94) or deep cells (n = 85). No significant differences between groups (t-test, p = 0.61, n = 3 birds) H. Scatter plot of *α* plotted across different depths. No correlation found between α and depth (Pearson’s correlation coefficient R = 0.05, n = 179 cells), with no significant linear relationship (p = 0.53).

Next we fit the displacement as a function of time using the formula: *MSD* = *βτ^α^* where *β* (diffusion coefficient) and *α* (scaling factor) are free parameters. The value of *α* distinguishes different forms of diffusion, such as random walk (*α* = 1), ballistic motion (*α* = 2), or superdiffusion (1 < *α* < 2)^30–32^ (**Figure 6A,B**). This diffusion model was a good fit for migration of individual cells (75.7% with R^2^ > 0.95) as well as the population average (R^2^ > 0.98 overall). We examined how *α* varied across trajectories from different animals and brain regions. All regions were well fit by *α* ranging from 1.4 to 1.6 in HVC and 1.3 to 1.8 across different regions (n = 8 birds, R^2^ > 0.99) (**Figure 6C,D**, **Table 1**).

**Table 1.**
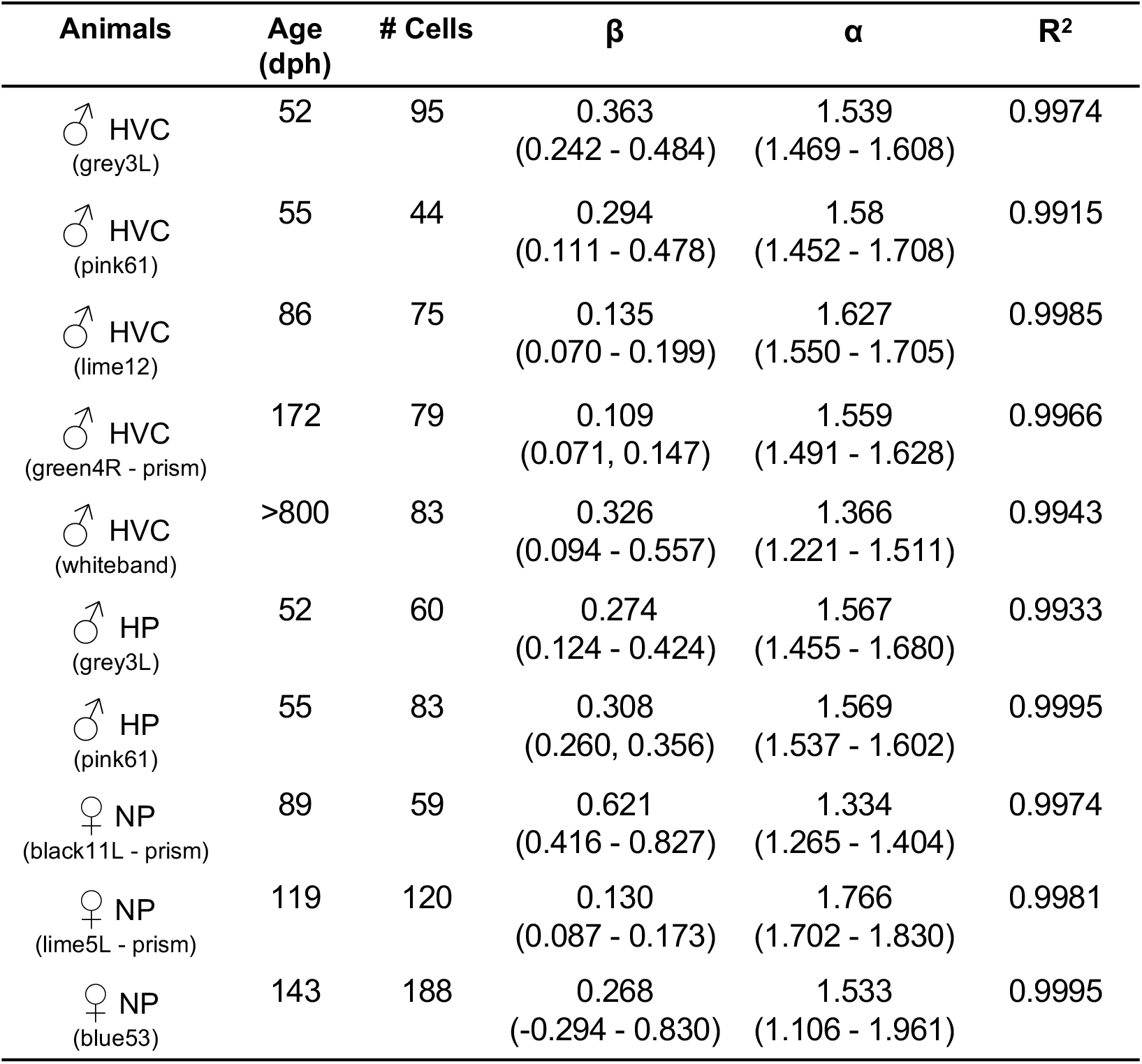
Diffusion model parameters across animals.

| **Animals** | **Age**<br>**(dph)** | **# Cells** | **$\beta$** | **$\alpha$** | **R<sup>2</sup>** |
| --- | --- | --- | --- | --- | --- |
| ♂ HVC<br>(grey3L) | 52 | 95 | 0.363<br>(0.242 - 0.484) | 1.539<br>(1.469 - 1.608) | 0.9974 |
| ♂ HVC<br>(pink61) | 55 | 44 | 0.294<br>(0.111 - 0.478) | 1.58<br>(1.452 - 1.708) | 0.9915 |
| ♂ HVC<br>(lime12) | 86 | 75 | 0.135<br>(0.070 - 0.199) | 1.627<br>(1.550 - 1.705) | 0.9985 |
| ♂ HVC<br>(green4R - prism) | 172 | 79 | 0.109<br>(0.071, 0.147) | 1.559<br>(1.491 - 1.628) | 0.9966 |
| ♂ HVC<br>(whiteband) | >800 | 83 | 0.326<br>(0.094 - 0.557) | 1.366<br>(1.221 - 1.511) | 0.9943 |
| ♂ HP<br>(grey3L) | 52 | 60 | 0.274<br>(0.124 - 0.424) | 1.567<br>(1.455 - 1.680) | 0.9933 |
| ♂ HP<br>(pink61) | 55 | 83 | 0.308<br>(0.260, 0.356) | 1.569<br>(1.537 - 1.602) | 0.9995 |
| ♀ NP<br>(black11L - prism) | 89 | 59 | 0.621<br>(0.416 - 0.827) | 1.334<br>(1.265 - 1.404) | 0.9974 |
| ♀ NP<br>(lime5L - prism) | 119 | 120 | 0.130<br>(0.087 - 0.173) | 1.766<br>(1.702 - 1.830) | 0.9981 |
| ♀ NP<br>(blue53) | 143 | 188 | 0.268<br>(-0.294 - 0.830) | 1.533<br>(1.106 - 1.961) | 0.9995 |

We fit the diffusion model to all individual cells tracked from these experiments (n = 876 trajectories, 8 birds) (**Figure 6E-F**). The distribution of *α* values from individual cells (n = 663 with R^2^ > 0.95 out of 876 trajectories) formed a unimodal distribution that had an average *α* = 1.77 +/- 0.5 (**Figure 6F**), suggesting that individual cells tended to behave in a superdiffusive manner across a population. Superficial cells (n = 94 cells) and deeper cells (n = 85 cells) did not differ significantly in their *α* parameters (two sample t-test, p = 0.61) (**Figure 6G**) and we found no significant relationship between the *α* of individual cells and the average depth of its tracked trajectory (using a linear regression test, p = 0.53, n = 179 cells), with no correlation with Pearson’s correlation coefficient (R = 0.05) (**Figure 6H**). Together, these results suggest that superdiffusion persists across different depths, brain regions, and ages, across both individual cells and the average population dynamics.

### Simulating neuron migration in a virtual songbird brain

Results thus far suggest that new neurons in the adult songbird HVC engage in a diffusion-like migration process. We designed a generative model of migration to evaluate whether this process is sufficient to allow neurons to disperse to integration targets in and around HVC over 3 - 4 weeks, which is the timescale for migration reported in previous studies^7^. In our model, simulated cells were born at random positions across a 2D plane representing the VZ (see **Methods**). After birth (at time *t* = 0) cells migrate throughout the brain in steps corresponding to 10 minutes of real world time. At each step, their position (P_t_) is updated by the following formula:

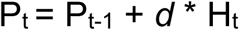

where P_t-1_ is its position at the preceding time point, *H_t_* is the cell’s heading for the current step, and *d* is the magnitude of the step being taken (**Figure 7A,B**). To reflect the changes in direction that cells exhibit during migration the cells heading, *H_t_*, is constructed using the following algorithms:

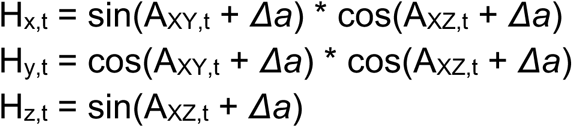

where A represents the angle direction that a cell is currently heading in XY or XZ and *Δa* is the change in heading angle (**Figure 7A**). The values of *d* and *Δa* are drawn stochastically at each time step *t* from distributions derived from experimental data (**Figure S5A,B**).

**Figure 7.**
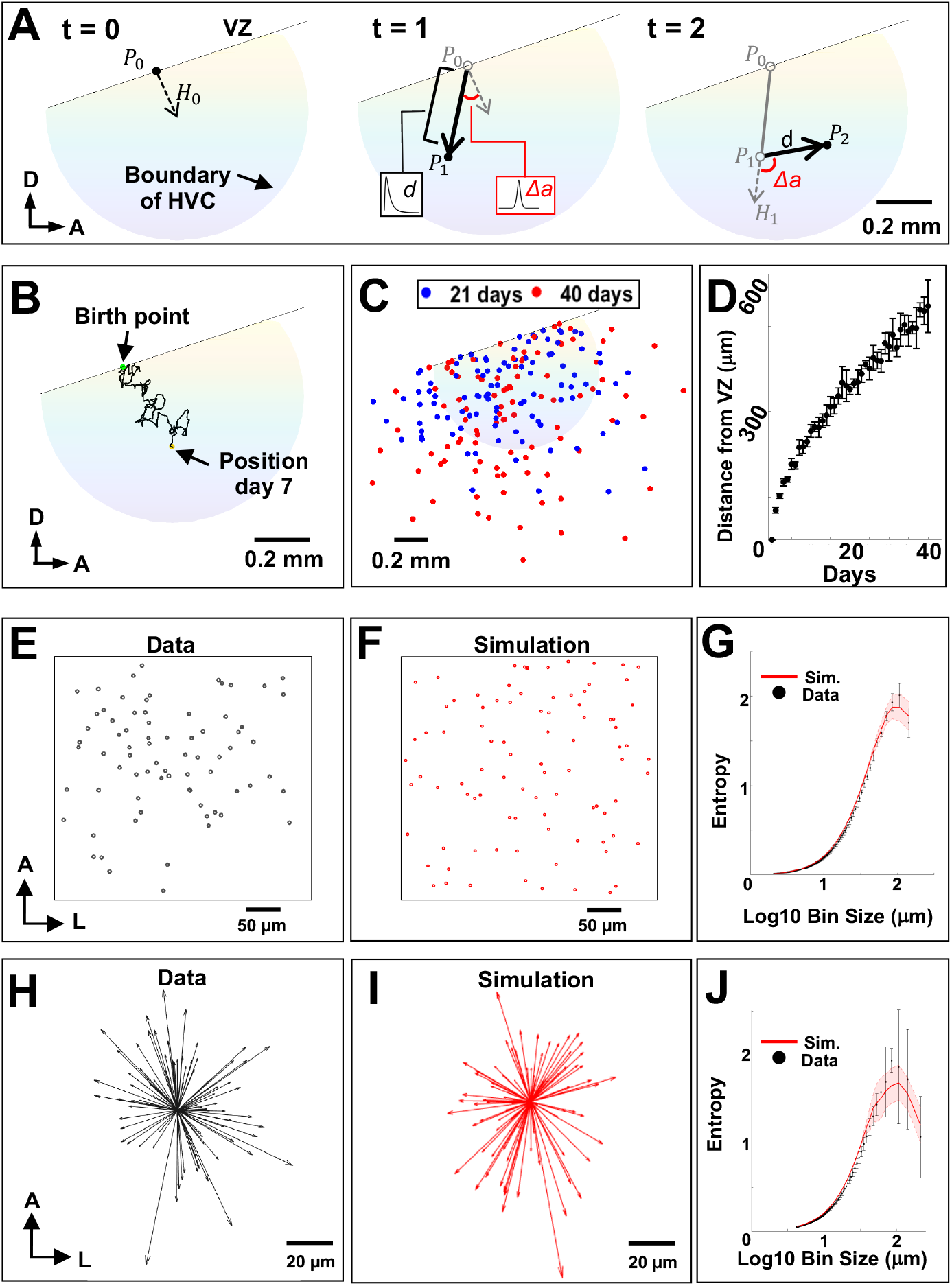
Simulating neuron migration in a virtual songbird brain. A. Schematic of movement rules for a simulated neuron within virtual HVC. *Left:* At t = 0, a cell is born at a random position (*P_0_*) with a heading *H_0_*; *Middle:* At t = 1, step size *d* and change in heading, *Δa,* are randomly drawn from probability distributions. The cell’s change in 3D position is calculated and the cell moves to its new position (*P_1_*) with its new heading; *Right:* This process repeats itself with *d* and *Δa* being drawn again and *P_t_* being calculated until the user-specified duration is complete. B. Example path of a simulated cell over 7 days of migration. The cell begins its migration from a point (green dot) on the VZ (black line) and travels for 7 days before reaching its current position (yellow dot). Cells move in 3D, but only movements in the sagittal plane are shown. C. Visualization of the positions of 100 simulated cells over 21 days (blue dots) and 40 days (red dots) of migration. D. Plot showing the average distance from the ventricle of 1000 simulated cells at each day of a 40 day simulation. Average distances of the simulated population (black dots) was recorded across 5 simulation runs (error bars). E. Positions of 96 migratory cells, simultaneously recorded *in vivo*, projected into a single plane. F. Positions of 96 simulated migratory cells after 21 days of migration projected into a single two-dimensional plane matching the dimensions of the imaging field-of-view shown in E. G. Entropy of cell positions from tracked cells (black dots) plotted alongside the entropy of simulated cell positions across time (red line) (n = 96 cells). H. Tracked cell headings in the horizontal plane, normalized to the same start positions (n = 96 cells). Headings are represented as vectors between the first and last recorded positions of the cell. I. Simulated cell headings in a two-dimensional plane, normalized to the same start positions (n = 96 cells). Headings were recorded after 21 days of migration. J. Entropy of cell headings from tracked cell headings (black dots) plotted alongside the entropy of simulated cell headings across time (red line) (n = 96 cells).

These simulated neurons exhibited similar migratory dynamics to cells observed *in vivo* (n = 143 cells across 2 birds, 3 hours of migration divided into 10 minute intervals). When measured across 3 hours, the paths of simulated cells had similar tortuosities compared with their *in vivo* counterparts and were well fit by a diffusive model with similar *α* parameters (µ_sim_ = 1.5539 +/- 0.0076, µ_track_ = 1.5595), indicating that the simulation was able to generate similar migration patterns compared with real cells (**Figure S5E-G,** see **Supplemental Methods**). Populations of simulated cells also exhibited a similar level of disorder compared to tracked cells. The entropy of both the position and headings of simulated cells was similar to their tracked counterparts (n = 96 cells, percent error_position_ = 4.69%, percent error_heading_ = 1.38%) (**Figure 7E-J**). Altogether, these results suggest that this simulation recapitulates several key features of new neuron migration in the postnatal zebra finch brain.

We next simulated the migration of 1000 new neurons in HVC over multiple weeks (**Figure 7C**). After 21 days, simulated cells were found throughout HVC and the surrounding tissue up to 1.580 +/- 0.065 mm from the VZ. The population center of mass (COM) was located 373.33 +/- 3.07 μm from the VZ, which is similar to the distance from the VZ to the center of HVC *in vivo* (374.35 μm)^33^. At the end of 40 days, simulated cells had dispersed further, migrating up to 2.16 +/- 0.345 mm from the VZ with a COM of 536.4 +/- 10.46 µm from the VZ (**Figure 7D**). These data suggest that this form of migration is sufficient to distribute cells throughout HVC and beyond over the biologically relevant timescale for migration and integration.

## Discussion

Here, we evaluated UBC-GFP zebra finches, a previously developed transgenic zebra finch line^12^, for the study of neuron migration. Using histological analysis, we found that GFP expression in the postnatal brain is primarily localized to mature and immature neurons. Grafting brain tissue from transgenics into wildtype birds revealed the presence of GFP+ cells that could migrate into the host brain and differentiate into neurons. Furthermore, using *in vivo* imaging, we found that GFP expression is strong enough and sparse enough to allow tracking of large populations of migratory cells in the intact brain.

Using these transgenic songbirds, we provide an in-depth quantitative characterization of neuron migration in the postnatal zebra finch brain. We found that migratory cells are present throughout HVC and surrounding tissue at high density. These cells dispersed in all directions and made frequent turns. We did not observe strong correlations between the speed or headings of adjacent cells, suggesting that neuroblasts may migrate independently. Migrating neurons were occasionally observed near vasculature for a portion of their trajectories but did not seem to rely on it exclusively as a scaffold. Finally, through computational analysis of the paths of hundreds of neurons, we found that this migration is well fit by a superdiffusive model and is sufficient to populate HVC *in silico*. Together these results reveal a form of superdiffusive migration that underlies new neuron dispersion in the juvenile and adult brain.

### Comparison with other forms of migration

Previous work in songbirds has predicted the existence of diffusion-like migration in and around HVC. Vellema and colleagues used mathematical modeling of the positions of newborn, BrdU-labeled cells to predict a ‘random migration’ that could explain patterns of cell dispersal in the canary brain^34^. This diffusion-like migration was also supported by *in vivo* imaging of retrovirally labeled newborn neurons in zebra finches in the anesthetized, juvenile HVC song nucleus^11^. In the present study, we provide additional evidence to support this prediction through detailed quantitative characteristics of hundreds of new neurons. We also show this form of diffusion-like migration extends beyond the boundaries of HVC and that it persists well into adulthood in both males and females.

Diffusion-like migration has also been observed in other neural systems, however not at the spatio-temporal scale we describe. For example, in the developing cortex, before, during, or after cells’ association with radial glia, new pyramidal neurons may exhibit diffusive-like movements^35–37^. It has also been observed in *ex vivo* time-lapse experiments that tangentially migrating interneurons undergo random-walk-like movements in the developing cortex, not dissimilar to the migration that we describe here^38,39^. Furthermore, *in vivo* time-lapse experiments showed that migratory interneurons with bipolar morphology move in many different directions in the intact developing cortex^40,41^. These tangentially migrating interneurons do not seem to be dependent on radial glial scaffolds but instead may be able to switch their migratory strategy between scaffold-associated and scaffold-free migration, especially once they arrive at the cortex^18,42–45^. Finally, in the adult rostral migratory stream (RMS), a major route for newborn neurons in the adult mammalian brain, timelapse experiments showed that almost a third of migration cells exhibited local, “exploratory” dynamics that changed direction frequently^46,47^.

However, there are key differences between these previously described forms of diffusive-like migration and what we describe here. The first key difference is that, in the developing cortex and the adult RMS, diffusive-like migratory dynamics are more restricted in space and time than in the songbirds. Among radially migrating projection neurons in the mammalian cortex the diffusive-like migration occurs at a brief phase of migration within the subventricular zone that lasts about 24 hours, prior to the resumption of radial migration^36,37,48^. In tangentially migrating neurons, diffusion-like dynamics are likely restricted to either the marginal zone or subventricular zone/intermediate zone layers, where ‘multidirectional migration’ occurs, before the postnatal invasion of the cortical plate radially^40,41,49,50^. Lastly, the migratory neurons of the RMS are spatially confined to a narrow band in the ventral forebrain, unless there is nearby tissue damage^19,51,52^.

A second key difference is that all these other streams of migration are highly directed. Radially migrating cells’ trajectories are oriented perpendicular to the VZ, tangentially migrating neurons migrate hundreds of micrometers parallel to the VZ before switching to radial migration in the cortex, and in the case of the RMS, migration is directed towards the olfactory bulb^48,51,53^. Thus, the diffusive-like patterns observed in time-lapse experiments from these systems appear to be relatively small deviations from an overall directed pattern of migration.

It is unclear why the form of neuron migration we observe differs from migration in these other systems. One possibility is that differences in experimental or analytical methods may accentuate the differences in migration patterns across these systems. Perhaps future quantitative analyses comparing large numbers of migratory neurons across these systems *in vivo* will reveal more similarities. Another possibility is that diffusion-like migration is more prominent in non-laminar structures, such as brain nuclei, and less prominent in laminar structures, such as the neocortex. Perhaps further analysis of neuron migration in the development of nuclear structure in the mammal will reveal more widespread diffusive-like patterns. Interestingly, recent work in amygdala tissue of adult humans discovered the existence of immature neuroblasts with embryonic origins that may also migrate in an undirected manner^54^. Finally, it is possible that the form of migration we observe is a specialization of migration in the adult avian brain to facilitate the replacement of lost cells in a flexible and efficient manner.

### Superdiffusive migration may offer efficient dispersal

Superdiffusion has been observed across a range of motile biological systems including intracellular molecular motors, super-spreading cancer cells, cerebellar neuroblasts, and predatory animals^26,55–57^. In these systems, superdiffusive dispersal dynamics have been proposed to be advantageous in sampling an environment with sparse cues. Whereas a random walk is prone to oversampling, and a more directed, ballistic strategy maximizes distance but undersamples, superdiffusion can be optimal^58–60^. For example, the superdiffusive searching behaviors of immunocyte T-cells may enable the immune system to find and destroy rare targets more efficiently^24^. At the organismal level, superdiffusivity may play a role when prey is sparse and may offer predators, such as sharks, a flexible strategy to meet environmental demands^57^. Although the mechanisms giving rise to superdiffusion are not yet clear, perhaps superdiffusive migratory neurons in adult songbird brains are more efficient at sampling their local environment and finding targets like migratory cues or prime integration targets within the circuit.

Although, mathematically, superdiffusion implies a stochastic process, it is unclear whether new neurons make flexible decisions as they migrate, or, alternatively, follow a predetermined but highly tortuous path. Within HVC, neurons were not observed to exclusively follow two well-described scaffolds in the embryonic and adult brain, namely radial fibers and vasculature. However, we cannot rule out the possibility that cells migrate along radial fibers outside of HVC or follow vimentin negative fibers.

### Potential mechanisms of neuron migration in songbirds

Although the precise mechanisms that guide neurons on their route through the zebra finch brain are unknown, there are several known molecular factors that control neuron migration in songbirds. For example, brain-derived neurotrophic factor (BDNF) is known to promote neuron migration in explant cultures from the canary brain^61^. In addition, the transition from expression of N-cadherin to NgCAM is thought to promote neuron migration in canaries^62^. Estrogen is thought to promote the departure of the cells from the VZ through an interaction with NgCAM in both zebra finch and canary explant cultures^63^. Studies *in vivo* show that the integration and survival of new neurons in HVC depends on testosterone, BDNF, and singing, as well as the death of adult neurons^3,64,65^.

In addition to molecular cues, the physical properties of the tissue, including stiffness, density, and topology, can influence cell migration^66–68^. For example, the shape of micropillars in a dish dictate whether migratory neurons will migrate directionally or undergo local, exploratory movements^68^. We speculate that these physical factors may be important to consider in the adult zebra finch as the mature brain poses several physical challenges for migrating neurons that are not present during development, such as increased cellular density, modifications of the extracellular space, increased vascularization, and the presence of highly stable functional structures, such as synapses^8,69^. Thus, there are many potential mechanisms, both molecular and structural, that may contribute to the superdiffusive dynamics observed here in the adult songbird brain.

### Simulation predicts existence of multiple forms of migration

Using our large dataset of hundreds of *in vivo* trajectories, we developed a simulation of neuronal migration. Our simulation revealed that diffusion-like migration is sufficient to populate HVC and nearby regions with new neurons across biologically relevant timescales. This generative model is part of a growing class of quantitative models that simulate cellular dynamics and pattern formation in tissue development^70–74^, including neuronal migration^75–78^.

Our simulation also makes several testable predictions. For one, our model predicts that cells are not predetermined to integrate in a particular region. Instead, their integration location is determined by a series of stochastic choices along their path. Additionally, in this model, cells are not influenced by environmental factors. For example, cell death or tissue damage would not direct migration towards lesioned areas. Finally, even after 40 days, stochastic dynamics alone were insufficient to reach the long distances (5 mm) described in previous experiments^7^. Others have proposed that migration could occur at different speeds in different parts of the avian brain, such as those enriched with radial glia^7,34^. This possibility motivates imaging studies in other brain regions.

### Experimental Outlook for UBC-GFP Songbirds

UBC-GFP zebra finches enable imaging of large populations of migrating neuroblasts *in vivo* without the need for viral labeling. The widespread expression of GFP could enable *in vivo* imaging in other regions of the brain in future studies with limited changes to the surgical preparation. Using 2PM through a cranial window we have been able to image migration 350 micrometers below the surface. However, three-photon microscopy can reach up to 1.3 millimeters *in vivo*^79^. This technology, in conjunction with UBC-GFP birds, could be used to image deeper regions of the zebra finch brain without the need for microprism implantation.

UBC-GFP songbirds could be also used to study the relationship between the birthplace of cells and their final integration site in the songbird brain. Previous studies in songbirds proposed that the birthplace along the VZ determines the specific cell identity and integration site of newborn neurons^4^. Brain grafting experiments between UBC-GFP birds and wild-type birds could be used to test this hypothesis.

## Conclusion

By applying *in vivo* 2P microscopy in transgenic zebra finches, we have quantitatively characterized a diffusion-like form of migration in the adult brain. Based on this data, we have developed a generative model that makes testable, quantitative predictions. We speculate that this migratory strategy enables neurons to efficiently navigate through the complex environment of the mature nervous system. Finally, our work demonstrates the value of UBC-GFP transgenic songbirds as a resource for future investigations of adult neurogenesis.

## Methods

### Animals

Animal use procedures were approved by the Boston University Institutional Animal Care and Use Committee (IACUC; Protocol #201800577) and carried out in accordance with National Institutes of Health standards. Birds were housed under standard conditions (12h/12h light/dark cycle, access to dry food and water ad libitum). Both males and females (50 - 800 dph) were used throughout the study. Birds were bred at Boston University. Wildtype mates were bred to heterozygote transgenics generating offspring of known age. Offspring were screened for GFP expression using a 440-460 nm blue illumination and 500 nm longpass glasses (Nightsea). Hatchlings exhibit green fluorescence throughout their body while in juveniles and adults, GFP expression is most readily observable around the eye.

### Surgical Procedures

Stereotaxic surgery was performed on a Kopf Instruments 940 digital frame with a 914 small bird adapter. Anesthesia was maintained with 1–2% isoflurane in oxygen. Male birds were injected with a pulled borosilicate capillary tube (Drummond Scientific) into AreaX (head angle 20°, 5.8 mm anterior, 1.5 mm lateral, and 2.8 mm below the dura) with 90 nL of DiI (D3911, Fisher Scientific), dissolved to 5 mg/mL with dimethylformamide (Sigma). Female and male birds underwent a 3 mm craniotomy over HVC coordinates as the centerpoint (relative to the bifurcation of the midsagittal sinus: head angle 35°, 0.3 mm anterior, 2.1 mm lateral). For cranial window experiments, a 3 mm diameter round coverglass (3 mm circular, #1, Thomas Scientific), bonded to a stainless steel cannula (304 S/S Tubing .125” OD x .115” ID x 0.019”; Ziggy’s Tubes and Wires, Inc., Sparta, TN) with UV-curing Optical Adhesive (#61, Norland), was placed over the exposed dura. For prism experiments, a custom implant was created by attaching a 1.5 mm micro prism with a reflective hypotenuse (Tower Optical) to a cranial window (described above) with optical adhesive. A microknife (FST) attached to the stereotaxic arm was used to cut the dura. Then, the prism implant was slowly lowered into the brain (10 μm/s) with the stereotaxic arm, using vacuum suction, until the cranial window was flush with the inner skull. The cannula walls were secured to the surrounding skull with cyanoacrylate adhesive. A custom kinematic headplate^14^ was secured to the skull with light-cured dental cement (Flow-It ALC, Pentron) and a curing LED 5W (NSKI, Amazon). The implant was covered with protective Kwik-Cast (VWR), which was removed before imaging. After recovery, finches were housed in a single cage in the colony room.

For allograft transplant experiments, transgenic zebra finches were anesthetized and underwent the craniotomy procedure over HVC (**Table S1**). Approximately 5-10 mg of tissue was dissected from HVC and the surrounding regions using a sterile biopsy punch (1.5 mm, Royaltek) and stored in sterilized EBSS (Worthington) on ice until implantation (< 30 min). A wild type host bird received a craniotomy and approximately 1 mm^3^ of tissue was removed via aspiration. Donor tissue was then gently fitted into the resulting cavity in the host brain. Gentle aspiration was applied peripherally to remove blood and saline. Surgery was concluded by replacing the host skull and sealing with silicone elastomer (Kwik-Sil, VWR) or by implanting a cranial window. Hosts were housed in a single cage within the colony for 3 - 9 weeks prior to sacrifice and histological characterization.

### Immunohistochemistry and Characterization of GFP expression

Zebra finches were euthanized with Euthasol (Virbac) and perfused with 30 mL of 1X phosphate-buffered saline (PBS), followed by 50 mL of 4% paraformaldehyde (PFA). Brains were extracted and stored in PFA overnight. After fixation, brains were washed with PBS and sliced into 50 μm sections with a vibratome (Leica; VT1000S).

The primary antibodies used were: a rabbit polyclonal anti-GFP in a 1:400 dilution (Millipore Sigma, AB3080, lot: 3717722), a mouse monoclonal anti-NeuN in a 1:500 dilution that is a marker for mature neurons (Millipore Sigma, MAB377, clone A60, lot: 3612227)^80^, a mouse monoclonal anti-DCX in a 1:400 dilution that is a marker for immature, migratory neurons (Santa Cruz Biotechnology, sc-271390 (E-6), lot: C3121)^81–83^, a mouse monoclonal anti-Vimentin (40E-C-s) in a 1:10 dilution that is a marker for radial glia^84^ (Developmental Studies Hybridoma Bank, AB528504), and a mouse monoclonal anti-HuC/HuD that is a marker for cells in neuronal lineage^85^ in 1:500 dilution (ThermoFisher Scientific A-21271). The secondary antibodies used were: Alexa Fluor 488 conjugated goat anti-rabbit (Abcam), Alexa Fluor 555 conjugated goat anti-mouse (Invitrogen).

Tissue sections were placed in a blocking solution composed of 2% nonfat dry milk (Great Value), 0.02% Triton X-100 (Sigma), and 1X PBS. Sections were incubated for 1 hour at room temperature in primary antibody diluted in blocking solution, washed 3 times with PBS, then incubated with secondary antibody diluted in blocking solution for 1 hour at room temperature or overnight at 4°C. Sections were mounted on microscope slides (Fisher Scientific) using Vectashield Antifade Mounting Medium with DAPI (Vector Laboratories, Inc.). Histological fluorescent images were acquired using a confocal microscope (Zeiss LSM800, Germany).

Image processing was performed with FIJI/ImageJ^86^. Individual ROIs were manually drawn around cell bodies in single Z-planes in one isolated channel. ROIs were then overlaid on the second channel for analysis, mean intensity in the GFP channel was measured, and ROIs were manually scored as colocalizing or not colocalizing with another antibody. Analysis was performed in Google Sheets or MATLAB.

### Fluorescent *in situ* hybridization chain reaction (HCR)

Fluorescent *in situ* hybridization chain reaction (HCR-FISH) was performed using custom probes designed by Molecular Instruments for detection of zebra finch GAD1, and UTS2B mRNA. Birds were perfused with formalin and brains were post-fixed in formalin overnight at 4°C. Brains were sectioned on a LEICA VT 1000S into 150μm slices and collected in 1X RNAse-free PBS. Slices were incubated in 90% DMSO/RNAse-free PBS for 2 hours at room temperature and then in 1% NaBH4/RNAse-free PBS for 15 minutes at room temperature. The slices were then incubated in 8% SDS/RNAse-free PBS for 1.5 hours at room temperature. Then, the slices were washed three times for 1 hour in 2X sodium chloride sodium citrate 0.1% Tween 20 (2X SSCT) at room temperature and then washed in probe hybridization buffer (Molecular Instruments) for 30 minutes at 37°C. The slices were hybridized overnight at 37°C in a solution containing 4 nM of each probe in hybridization buffer. On the second day, the slices were washed four times in wash buffer (Molecular Instruments) at 37°C and three times in 5X SSCT for 5 minutes at room temperature. Then they were incubated in an amplification buffer for 30 minutes at room temperature. Lastly, they were incubated overnight in a solution containing 2μL of fluorescent-carrying hairpins per 200μL of solution (provided by Molecular Instruments) at room temperature. The next day, slices were washed three times in 5X SSCT and one time in 5X SSC at room temperature. They were then fixed for 20 minutes in PFA 4% at room temperature. Subsequently, immunohistochemistry staining was performed using the following antibodies: rabbit polyclonal antibody against GFP (Millipore Sigma, AB3080, lot: 3717722) and Alexa Fluor 488 conjugated goat anti-rabbit (Abcam). All images were acquired using a confocal microscope (Zeiss LSM800, Germany) and loaded into FIJI/ImageJ for quantitative HCR-FISH staining analysis in the manner described above.

### Two-photon Microscopy

Imaging was performed on a commercial 2P microscope system (Bruker Corporation, Prairie View Software). GFP, sulforhodamine-101, and DiI were excited by near-infrared light (920 nm) produced by a titanium:sapphire laser (Spectra-Physics, Insight X3; or Coherent, Chameleon Ultra II). Post objective power was less than 100 mW for all imaging experiments. Images were acquired using a Nikon 16X, 0.80 NA water-immersion objective lens (ThorLabs, N16XLWD-PF), or an Olympus 20X, 0.50 NA water-immersion objective lens (Olympus, UMPLFLN20XW), and photomultiplier tubes (Hamamatsu, H10770PB-40).

Imaging was performed in head-fixed and body restrained unanesthetized animals. Headplates were attached magnetically to a custom kinematic fixation system^14^ on top of a goniometric stage (ThorLabs) while the animal was restrained in a custom foam restraint. A custom 3D-printed conical well was fastened to the headplate with silicon elastomer to hold water for longer timelapse sessions. The animal was observed with an infrared camera (USB100W05MT-DL36, ELP, iSpy Software). Time intervals varied per animal (**Table S2**). In some sessions the Z-stacks were sampled in succession (∼10 minute interval between frames) for up to 3 hours. In longer sessions, timelapses were acquired intermittently at intervals ranging from 20 minutes to up to an hour between Z-stacks for 5 -12 hours. Between imaging timepoints, the animal was freed from restraint and returned to its cage with food and water. In some sessions, the animal was injected IM with 50 μl of 20 mM SR-101 (Invitrogen) before the last time point to label the blood vessels in the volume.

### Image Processing and Data Analysis

All data was stored and accessed on Boston University’s Shared Computing Cluster. Session images were loaded into FIJI as 4D hyperstacks. A denoising algorithm was applied to the hyperstacks using FIJI functions in the following order: Background Subtraction, a Gaussian high pass filter, Median Filter, Despeckle. Hyperstacks were registered with the Correct 3D Drift plugzin in FIJI. The session data was first split up by brain region, and then split up into four quadrants for easier distribution to trackers and management for tracking. The centerpoint of migratory cells was identified by expert trackers across time using a custom macro script in FIJI that enabled the use of Orthogonal Views and sequential region-of-interest (ROI) selection with the Multi-Point Tool.

All track analyses were performed using custom scripts in MATLAB. Tracking data was pre-processed to remove duplicate tracks and erroneous points. Cells that traveled less than 5 microns and cells with tortuosities above 7 were removed.

To evaluate the degree of “association” of migrating cells with vasculature, tracks from imaging sessions with SR-101-labeled blood vessels were evaluated in FIJI. After the GFP channel was isolated and migratory GFP+ cells were tracked by expert trackers, GFP and SR-101 channels were recombined. Expert scorers overlaid tracked ROIs (the centerpoint of each cell across time) and recorded the position at that time point as an “association” if the ROI came within 1 micron of SR-101-positive vasculature. Length of associations were determined by computing the distance traveled by cells that had ROIs closely associated with SR-101 labeled vasculature for multiple timepoints consecutively.

Custom scripts in MATLAB were used to fit a power law model to either the MSD of individual cells or the average MSD from entire cell populations to extract the *α* parameter. This was done using the MATLAB *fit* function, with ‘power1’ specified as the model. *α* fit to individual cells was only considered if it was less than 4 and the R^2^ value of the fit was above 0.95.

### Migration simulation

The migration simulation was written in MATLAB. HVC was represented as a 1 mm diameter sphere intersected by a tilted plane (representing the VZ) and has a comparable volume and shape as HVC found in zebra finches^33,87^.

A simulation run began with the birth of a cell in the VZ at a randomly selected point on the VZ-HVC interface. Each cell (represented as a vector with position *P* and heading *H*) was born heading in a random 3D direction which serves as the cell’s initial *H*. At each timepoint (whose real world equivalent duration is 10 minutes) a cell takes a step of distance *d* in the direction *H_t_*. *d* is drawn from a log-normal distribution (μ = 0.83 μm, σ = 0.79 μm) and *Δa* is drawn from a logistic distribution (μ = -1.38°, σ = 9.79°, p = 0.9) with a uniform offset (ranging from -180° to 180°, p = 0.1; **Figure 7A**). These distributions were derived from step sizes and angle changes measured from *in vivo* cell migration data recorded at ∼10 minute intervals (see **Supplemental Methods**).

Cells are prevented from crossing the VZ at any point during their migration: if P_t_ is on the other side of the VZ, *d* and *Δa* for that step are redrawn.

## Acknowledgements

We thank Timothy Otchy, Mimi Kao, Sinead O’Brien, Amber Hickey and the Boston University Animal Science Center for support and maintenance of the finch colony. Jack Giblin, Martin Thunemann, Natalie Thunemann, Ryan Senne, Su Jin Kim, Amy Monasterio, Steve Ramirez, Noshin Nawar, Jeroen Eyckmans, Todd Blute, and Boston University Research and Computing Services provided equipment and technical support. We thank Young Ye for tracking assistance and help with the tracking pipeline. Peien Lyu, Christa Rose, Margaret Seo, and Jacqueline Wang for tracking assistance. Gabe Ocker, Lei Tian, and Yujia Xue for useful conversations about data analysis. Naomi Shvedov was supported by an NSF NRT fellowship. This project was supported by seed funding from the Boston University Neurophotonics Center.

## Author Contributions

Conceptualization: N.R.S. and B.B.S.; resources: T.J.G.; data collection: N.R.S., T.D., S.A., B.L.B; modeling and analysis: N.R.S. and S.A.; writing – original draft: N.R.S; writing – review & editing: N.R.S., S.A., T.D. and B.B.S.

Authors have no conflicts of interest to declare.

