## Supplemental Materials for "In vivo imaging in transgenic songbirds reveals superdiffusive neuron migration in the adult brain"

Contents

**Supplemental Figures** **2**
Figure S1. GFP expression varies across brain regions and cell types 2
Figure S2. Comparison of trajectory estimates in the imaging and axial dimensions 3
Figure S3. Similar dispersal dynamics are observed across brain depths 4
Figure S4. Interactions between migratory neurons and vasculature 5
Figure S5. Simulation recapitulates statistics of experimental data 6
Figure S6. Data from birds with cranial windows and prisms. 7
**Supplemental Tables** **8**
Table S1. Quantification of transplanted cells across different hosts 8
Table S2. Imaging information for all birds 9
**Supplemental Methods and Results** **10**
Identification of UBC-GFP transgene insertion site 10
Validation of migration simulation against experimental data 10
Simulation extrapolation analysis 11
Estimation of angle and step size distributions 12
Calculating entropy 12
**Supplemental Videos** **14**
**Supplemental References** **15**

### Supplemental Figures

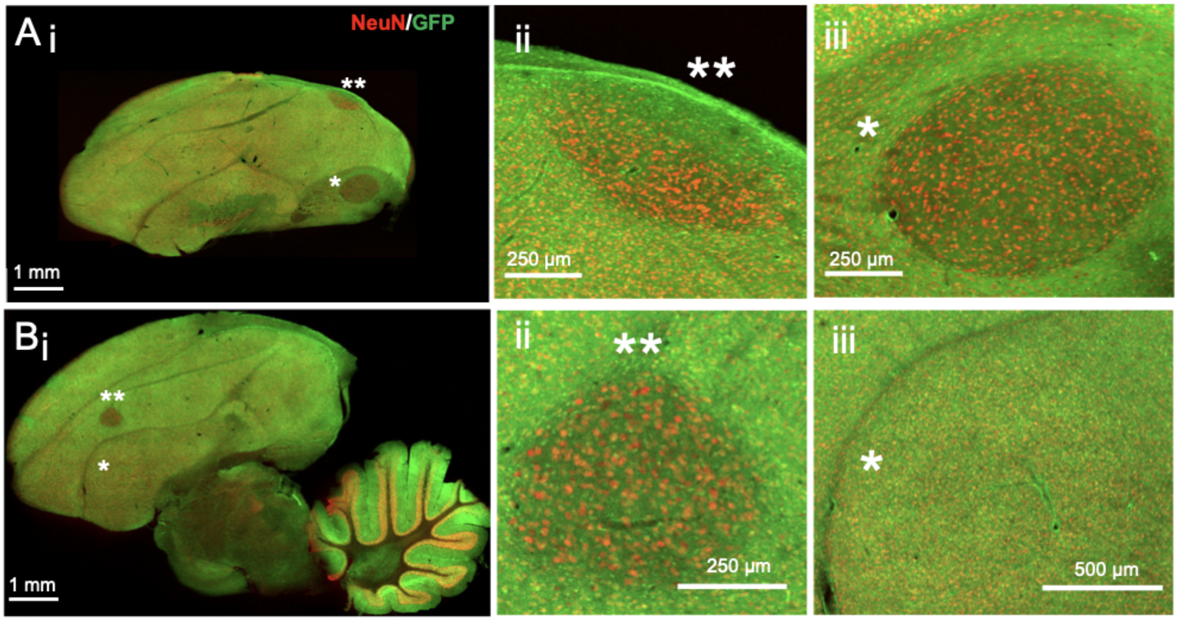


Figure S1. GFP expression varies across brain regions and cell types

A.(i) Transgenic GFP+ brain slice showing HVC (ii, 2 asterisks) and RA (iii, 1 asterisk) co-labeled with NeuN and GFP. GFP is dimmer in HVC and RA, regions that see relatively fewer to no neuron addition. B.(i) Transgenic GFP+ brain slice showing LMAN (ii, 2 asterisks) and Area X (iii, 1 asterisk) co-labeled with NeuN and GFP. GFP is dimmer in LMAN, a region with no neuron addition.

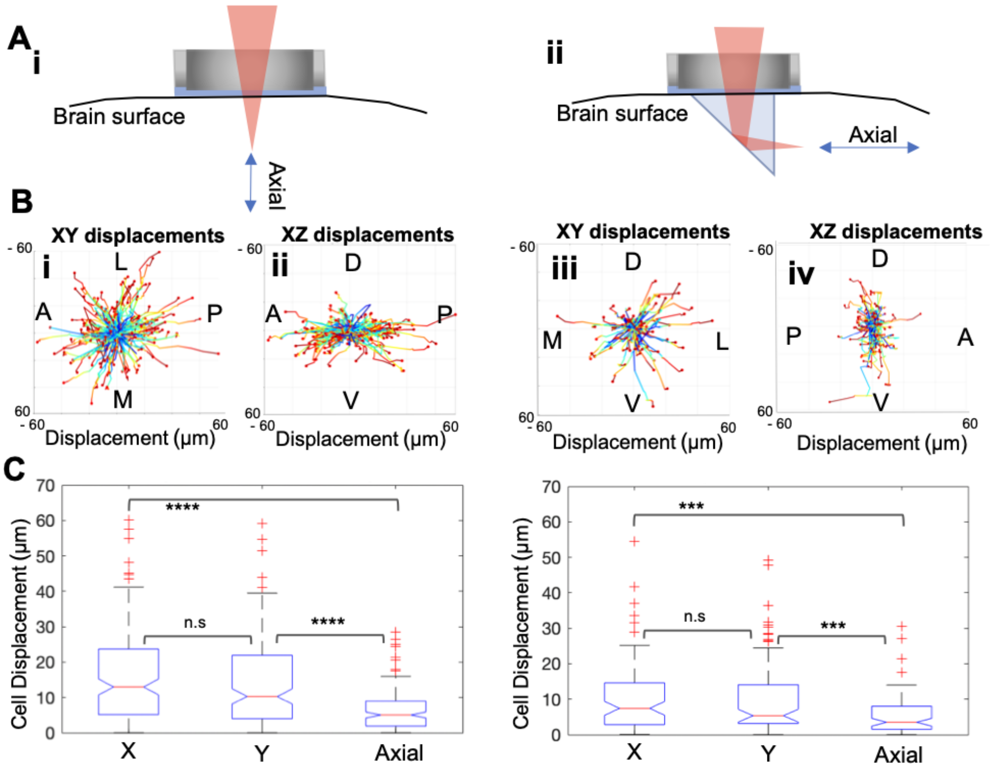


Figure S2. Comparison of trajectory estimates in the imaging and axial dimensions

A.(i) Schematic of axial dimension through cranial window implant and (ii) Schematic of axial dimension through microprism implant. B.(i) “in plane” relative XY displacements along the AP/ML axes for cranial window and (ii) “out of plane,” axial, relative XZ displacements along the AP/DV axis for cranial window (n = 169 migratory cells, 1 bird, 4.5 hour imaging session); (iii) for prism placed with imaging face oriented DV and ML, in plane, relative XY displacements are shown, and (iv) in prism, out of plane, relative AP/DV displacements are shown (n = 88 migratory cells, 1 bird, 3.7 hour imaging session). C. *Left:* Cranial window notched box plots. “In plane” (XY) displacements are not significantly different from each other (two-sample t-test, p = 0.13) but the axial dimension is significantly lower than both X and Y (two-sample t-test, p = 1.5e-16, and p = 1.9e-12, respectively). *Right:* Prism notched box plots. “In plane displacements” (XY) do not differ significantly from each other (two-sample t-test, p = 0.92), while the axial dimension is significantly lower than both X and Y (two-sample t-test, p < 0.0001 for both).

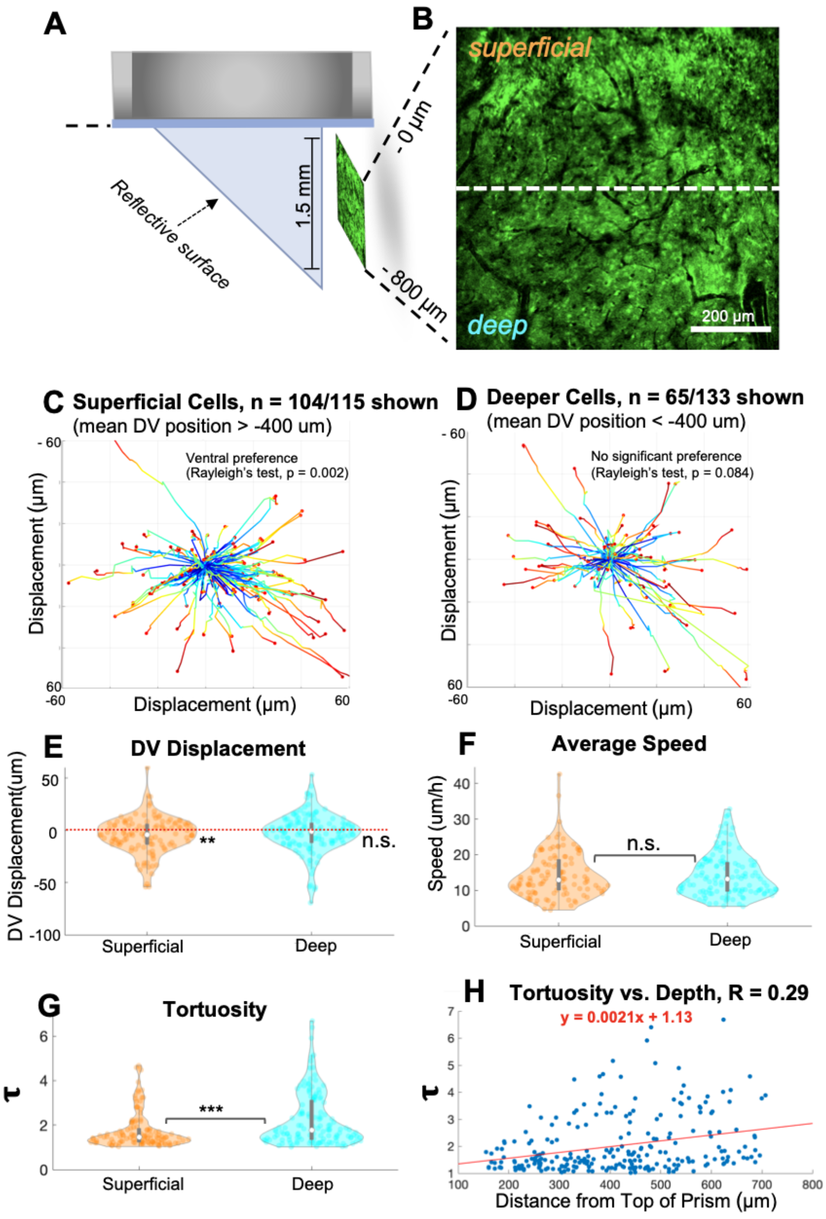


Figure S3. Similar dispersal dynamics are observed across brain depths

A. Microprism implant assembly. B. 2P fluorescence imaging FOV. C. Representative sagittal plane DV displacement vectors shown from 104 superficial tracks across 2 prism birds. D. Representative coronal plane DV displacement vectors shown from 65 deeper tracks in the same 2 birds. E. Slight ventral bias detected in superficial cells (1.22 µm/h ventrally; t-test, $p = 0.0043$; n = 115 cells, 3 birds). F. No difference in speed between superficial (n = 115) and deeper (n = 133) groupings of tracks (t-test, $p = 0.92$). G. Deeper tortuosities (n = 133) are significantly higher than superficial tortuosities (n = 115) (two-sample t-test, $p = 0.0003$). H. A positive correlation is found between tortuosity and depth (Pearson's correlation coefficient $R = 0.29$) with a significant linear relationship ($p = 3.4e\text{-}6$).

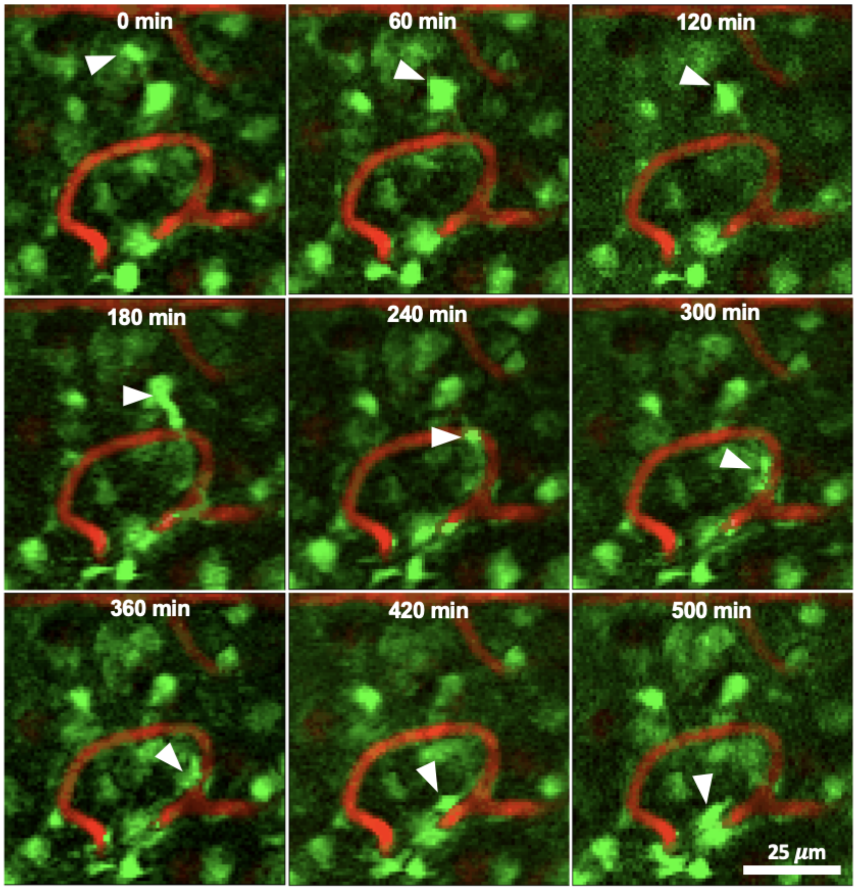


Figure S4. Interactions between migratory neurons and vasculature

Example GFP+ migratory cell coming in vicinity of SR-101-labeled vasculature and changing migratory direction during interaction. These images are standard deviation depth projections across 50 microns and imaged from an 86 dph male bird HVC across 8.3 hours.

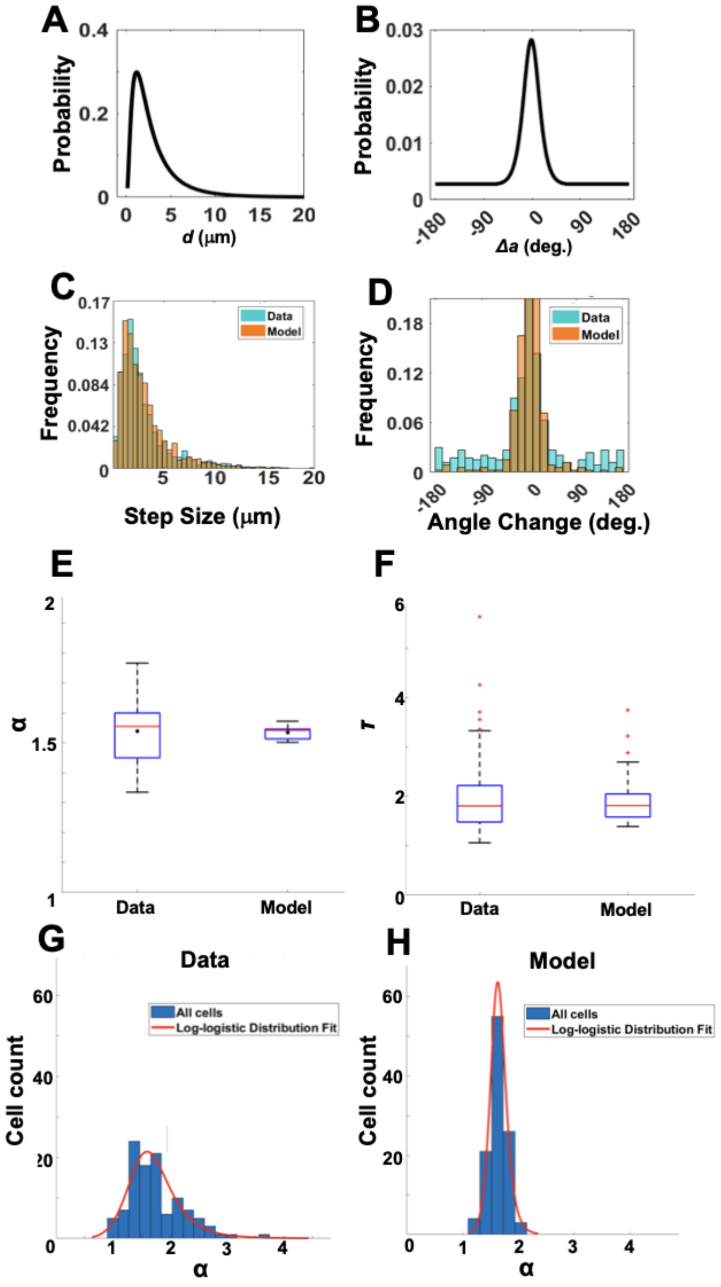


Figure S5. Simulation recapitulates statistics of experimental data

A. Lognormal probability distribution of *d* used in this simulation (μ = 0.83, σ = 0.79). n = 2393 steps from 143 cells across HVC of male birds aged 52 - 55 dph. B.Probability distribution of *Δa*. The distribution is the weighted sum of a logistic (μ = -1.38, σ = 9.79, p = 0.9) and a uniform random distribution offset (ranging from -180° to 180° with p = 0.1). n = 2393 steps from 143 cells across HVC of male birds aged 52 - 55 dph. C. Distribution of step sizes recorded from experiments overlaid with simulation step sizes. D. Distribution of angle changes recorded from experiments overlaid with the distribution of angle changes recorded from simulation. E. Measured population *α* values from tracking experiments alongside simulation model runs. Black dots represent the mean population *α* ($\mu_{track}$ = 1.5385, $\mu_{sim}$ = 1.5338). F. Mean tortuosity values from in vivo tracking experiments alongside the simulation model. G. Histogram of *α* parameter for individual tracked cells (n = 110 cells, 2 birds). The data is well fit by a unimodal log-logistic distribution. The average *α* of good fits ($R^2$ > 0.95, *α* < 4 ) is 1.7457 +/- 0.4798. H. Histogram of *α* parameter for individual simulated cells (n = 110 cells). The data is well fit by a unimodal log-logistic distribution. The average *α* of good fits ($R^2$ > 0.95, *α* < 4 ) is 1.6414 +/- 0.1609.

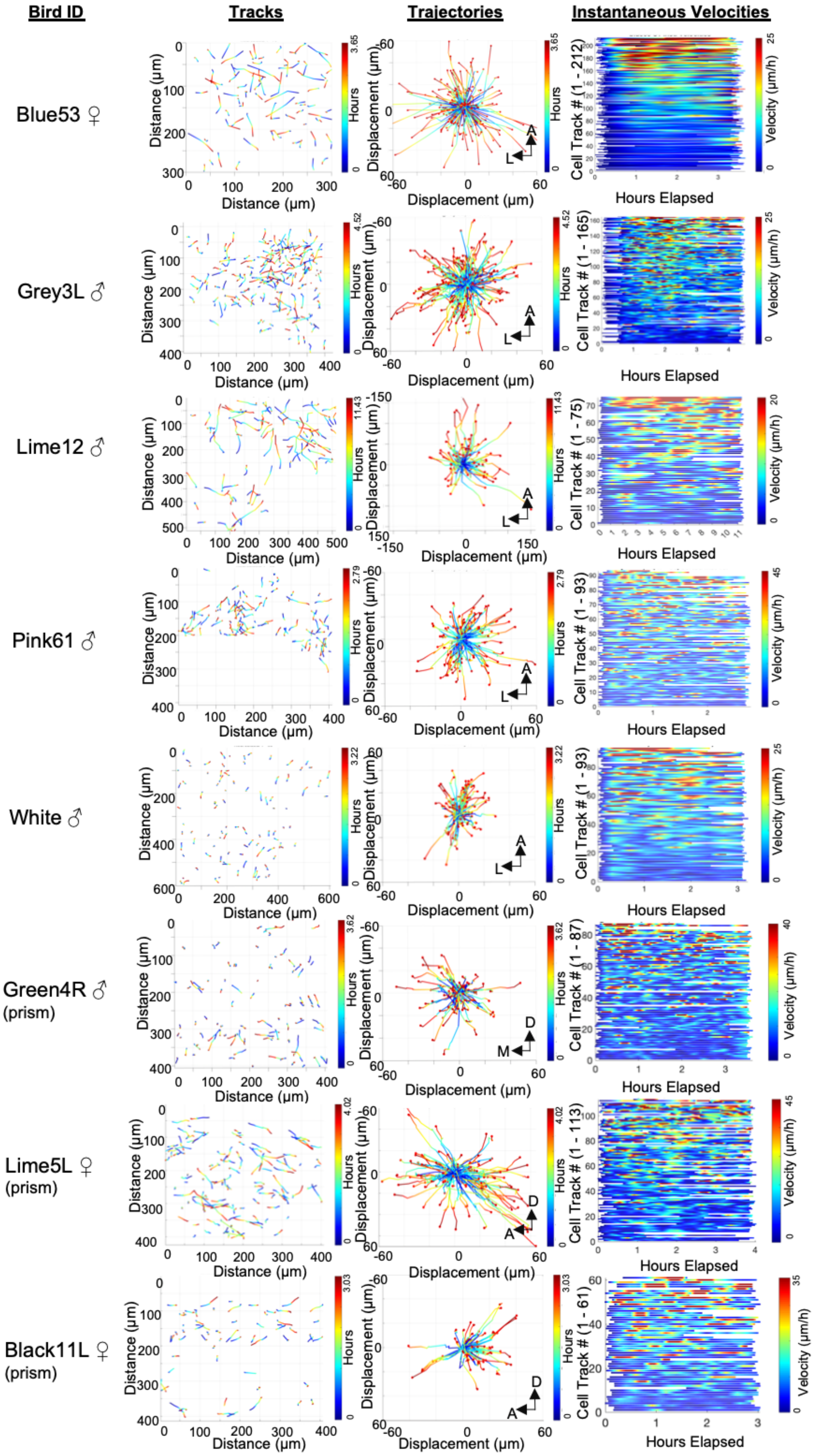


Figure S6. Data from birds with cranial windows and prisms.

Summary imaging data from 8 birds, showing trajectories, displacements and instantaneous velocities. See **Table S2** for imaging details.

### Supplemental Tables

Table S1. Quantification of transplanted cells across different hosts

| Donor (dph/ Sex) | Host (dph/ Sex) | Days post transplant | # Cells Observed | # Slices Examined (50 μm thick) | Mean distance to graft (μm) | Max distance from graft (μm) |
|---|---|---|---|---|---|---|
| 55/F | >730/F | 21 | 21 | 19 | 124.00 | 350.30 |
| 49/F | 113/F | 48 | 2 | 38 | 165.21 | 165.21 |
| 54/F | 80/F | 34 | 3 | 37 | 1342.70 | 1943.534 |
| 54/F | 101/F | 34 | 23 | 20 | 1010.35 | 2018.504 |
| 53/M | 97/M | 61 | 1 | 41 | 188.126 | 188.126 |
| 53/M | 120/M | 61 | 6 | 35 | 334.302 | 761.778 |

Table S2. Imaging information for all birds

| Bird ID | Implant | Imaging Age (dph) | Sex | # of Sessions | Average Imaging Interval (min) | Session Duration (min) | # Cells Tracked |
|---|---|---|---|---|---|---|---|
| grey3L | Window | 52 | M | 1 | 13.7 | 271 | 155 |
| pink61 | Window | 55 | M | 2 | 9.80 | 167 | 127 |
| lime12 | Window | 86 | M | 1 | 59.8 | 686 | 75 |
| green4R | Prism | 172 | M | 1 | 9.62 | 217 | 79 |
| white | Window | >800 | M | 1 | 20.3 | 193 | 83 |
| black11L | Prism | 89 | F | 1 | 12.2 | 182 | 59 |
| lime5L | Prism | 119 | F | 1 | 12.1 | 241 | 120 |
| blue53 | Window | 143 | F | 1 | 48.9 | 219 | 188 |

### Supplemental Methods and Results

#### Identification of UBC-GFP transgene insertion site

UBC-GFP songbirds were previously created using lentiviral mediated trangenesis[1]. To identify the transgene insertion site we used whole genome sequencing followed by alignment to the zebra finch genome (bTaeGut2.pat.W.v2, https://www.ncbi.nlm.nih.gov/data-hub/genome/GCF_008822105.2/) and plasmid FUGW (Addgene plasmid # 14883 ; http://n2t.net/addgene:14883 ; RRID:Addgene_14883).

Genomic DNA was obtained from blood. The medial metatarsal vein in the leg of a male GFP+ zebra finch was nicked with a sterile 20 gauge needle. Blood was drawn with a glass Micro-Hematocrit capillary tube (Thomas Scientific). DNA was then extracted from 10 µl of blood using a genomic DNA extraction and purification kit (New England Biolabs). The sample was sent overnight on dry ice to a commercial next-generation sequencing company (LC Sciences, Houston, TX) for genome sequencing. The following processing and analysis steps were performed by LC Sciences: 1) Adapters and low quality reads were removed by Cutadapt and FastP softwares, respectively. 2) Data quality control was performed using the FastQC software. For genome alignment, BWA MEM[2] software was used. 3) Alignment of sequenced genomes was performed to determine the insertion site of the transgene. 4) Genome alignment to plasmid and reference genome was viewed in IGV, and allowed for insertion sites to be determined.

Through whole genome sequencing, we found that the transgene was inserted on Chromosome 9. The insertion occurred in the range spanning positions NC_045008.117066610 - NC_045008.1:17066705, which is found within a gene called von Willebrand factor A domain containing 5B2 isolate (VWA5B2). VWA5B2 is preferentially expressed in the adrenal glands, and throughout the brain[3]. The nearest gene to the insert, preceding VWA5B2 at the 5' end, was Dishevelled Segment Polarity Protein 3 (DVL3). The DVL3 protein is involved in the Wnt signaling pathway[4]. The nearest gene succeeding the VWA5B2 insert at the 3' end is alpha-1,3-mannosyltransferase (ALG3).

#### Validation of migration simulation against experimental data

In order to validate the design of the simulation, three main metrics were recorded to compare simulation output with cell tracking data: speed, tortuosity, and the α parameter derived from the curve fit to the MSD across time intervals. Each of these parameters were measured per cell for the entirety of their migration. These metrics were gathered from simulations that spanned a nearly identical time interval and window as that of cell tracking experiments (10 minute time interval, 3 hour tracking window) in order to be able to draw direct comparisons between the outcomes of the simulation and tracking data. A two sample t-test revealed that there was no significant difference in the tortuosity measured between the simulation and the tracking experiments, suggesting that the simulation was successful in recapitulating tortuosity dynamics that were observed in vivo (n = 143 cells, $p_{tort} = 0.0967$) (**Figure S5E**). Distributions of well-fit ($R^2 > 0.95$) individual $\alpha$'s measured for each trajectory were well-described by log-logistic distributions for both the simulated and tracked cell populations (n = 110 cells, $\mu_{sim} = 0.50$, $\sigma_{sim} = 0.055$, $\chi2_{sim} = 7.35$, $p_{sim} = 0.12$, $\mu_{track} = 0.51$, $\sigma_{track} = 0.15$, $\chi2_{track} = 9.41$, $p_{track} = 0.052$ (**Figure S5G-H**). Although both had a similar distribution peak, there was a noted difference in σ, indicating that although the average α from the simulation matches the observed α from in vivo, the population variability of $\alpha$ is not entirely recapitulated (**Figure S5E**). It should be noted that the average population $\alpha$ measured across 10 simulation runs is 1.535 (n = 10,000 cells , $\sigma$ = 0.0116), which defines the movement of simulated cells as superdiffusive. This superdiffusive mean population $\alpha$ is nearly identical to the mean population $\alpha$ recorded across 8 birds during tracking experiments ($\mu_{track}$ = 1.5385) (**Table 1**). For speed, a two sample t-test indicated a significant difference ($p_{speed} \cong 0$) between the simulation and tracking experiments although there is overlap between the two groups' standard deviations. However, the distributions of step size from the tracking experiments and the simulation are highly similar (paired sample t-test, p = 0.5341) (**Figure S5C**), suggesting that the difference in measured speed originates either from the simulation's assumption of independent stochastic step and heading selection, or the frequent migratory pauses cells take *in vivo* that are not accounted for in the simulation (see **Figure S6**).

Simulated cells in these validation experiments began their migrations at random points throughout HVC, not in the VZ, as tracked cells were distributed throughout the volume at the time of their recording. Speed was calculated as the sum of all distances traveled divided by the duration of migration. Tortuosity was measured as the ratio between the length of the path traveled by a cell to the shortest distance between its start and end points. Log-logistic curves were fitted to the distributions of individual cells' $\alpha$ using MATLAB's custom distribution fitter. In addition to calculating the individual $\alpha$ for each cell's migration, the population $\alpha$ was assembled from every cell simulated within a single run by taking the average of the MSD at each $\tau$ across all cells to form a single MSD curve that was representative of the simulated population.

##### Simulation extrapolation analysis

Once this model had been validated against the tracking data over a 3 hour time window, the simulation was run for 40 days to observe if superdiffusive dynamics could be successfully extrapolated to biologically relevant timescales. For these experiments, cells were born in the VZ to observe their behavior and spread for the entirety of their lifespan. To assess whether HVC was sufficiently populated, we took the average of each cells' minimum distance from the VZ across 5 simulation runs with each run containing 1000 cells. This distance was calculated and compared to the center of mass of the virtual HVC: for **Figure 7D**, the minimum distance was recorded per cell at the end of each day of migration. A similar process was used to calculate the average maximum distance of each cell from the VZ over 40 days to see how far cells could migrate away from their birthplace. Additionally, the Shannon entropy (*see Calculating entropy* below) of the final positions and headings of 1000 cells simulated over 21 days were compared to entropy of the positions and headings of tracked cells at the end of 3 hours of tracking. Simulated endpoints were recorded from cells that fell within the FOV used during cell tracking at the end of each day of migration; likewise, simulated cell headings were calculated as the vector between the cells final position and its positions 3 hours prior to its migration ending when both its final and 3 hour prior position fell within the aforementioned FOV. This FOV, which was a 3D volume (350 x 400 x 300 μm), was conserved from parameters used during cell tracking experiments and permits for direct comparisons to be drawn between the simulation output and the tracking data.

#### Estimation of angle and step size distributions

##### Angle

The $\Delta a$ distribution was obtained by fitting a logistic distribution to the histogram of XY plane angle change ($\Delta a_{XY}$) measured from tracking individual cells in vivo during cranial window 2P imaging conducted in transgenic finches wherein 143 cells were tracked over approximately 3 hours across two birds - these birds and these specific cells were purposefully selected from all experimental birds/tracked cells to form this distribution as these were juvenile male birds with cells recorded in HVC (pink61 and grey3L). Artifacts resulting from the manual cell tracking process were reduced by excluding recorded $\Delta a_{XY}$ when a cell migrated less than 5 μm before or after the $\Delta a_{XY}$. Based on prism tracking experiments that indicate that cells migrate similarly in both XY and XZ (see **Figure S2**), the $\Delta a_{XZ}$ distribution was inferred to be identical to $\Delta a_{XY}$. A logistic distribution was determined to be the distribution of best fit to the $\Delta a$ data based on a maximum log likelihood estimation. In order to account for rarer, but observed cases in which cells made broad (>90°) turns between steps, a secondary uniform distribution (μ = 0°, σ = 103.92°) ranging from -180° to 180° was included in creating $\Delta a$. At the beginning of each step, $\Delta a_{XY}$ and $\Delta a_{XZ}$ are independently and randomly generated for a cell. When either of these $\Delta a$ are generated, it will randomly select this angle change from the primary logistic distribution with a 90% probability, and from the secondary uniform distribution with a 10% probability.

##### Step size

The distance *d* that a cell will traverse over a single simulated step is randomly selected from a continuous lognormal distribution fit to the distribution of step distances from the aforementioned 2P cell tracking data.

#### Calculating entropy

The Shannon's entropy of the final recorded positions of the tracked cells and of the final positions of the simulated cells was calculated as follows:

$$Entropy = -\sum p(C_b) * log(p(C_b))$$

where $p(C_b)$ is the normalized probability that a cell's position is within any spatial bin when bin size is gradually increased. For each iteration of entropy calculated, the number of bins was increased by one, beginning with 3 bins and iterating until 200 bins. The number of cells within each bin is counted and the probability that a cell is in any of the bins, $p(C_b)$, is calculated and normalized to the total number of cells present in all bins. Inserting this probability into Shannon equation allows us to calculate the average entropy of the cells across the entire FOV. A similar approach was utilized when calculating the heading entropy of both tracked and simulated cells with heading being considered instead of the cell's final recorded position.

To create a maximum entropy model simulation, we performed the same procedure as for entropy calculations but assumed a multinomial uniform distribution by setting a constant probability of cells being in a bin.

#### Supplemental Videos

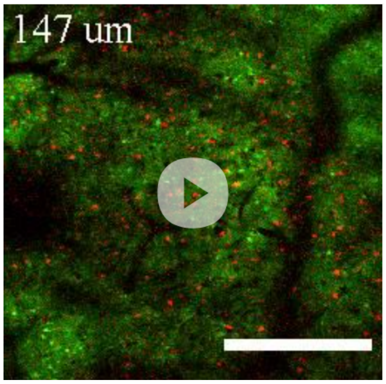


**Video S1. Z-stack in transgenic male zebra finch HVC**
Z-stack spanning the dura, hyperpallium, ventricular zone, and HVC of juvenile transgenic male. $HVC_X$ projecting cells are retrogradely labeled with DiI. Scale bar is 200 microns.

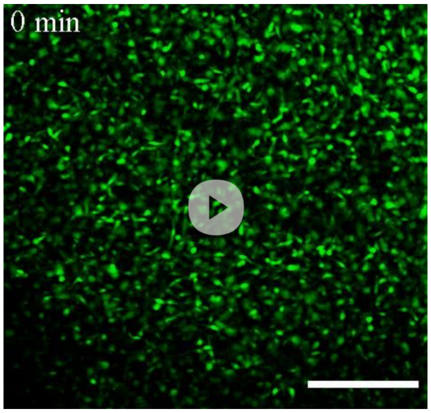


**Video S2. Large FOV time-lapse of migratory population**
Time-lapse 2P imaging of mature and migratory neurons in the adult female nidopallium. Movie is a standard deviation depth projection of 80 microns, background subtracted. Scale bar 100 microns. Movie loops four times at 5 fps.
